# Influenza virus Non Structural protein 1 (NS1) Chaperones Nucleoprotein (NP) Oligomerization to Coordinate Ribonucleoprotein Assembly and Genome Replication

**DOI:** 10.64898/2026.05.16.725629

**Authors:** Sayan Das, Aratrika De, Prabuddha Bhattacharya, Ayush Munshi, Puja Kumari, Dibyendu Samanta, Arindam Mondal

## Abstract

Influenza virus nucleoprotein (NP) co-transcriptionally enwraps viral genomic/antigenomic RNAs to assemble viral ribonucleoprotein complexes (RNPs), while the non-structural protein-1 (NS1) is classically recognized as an antagonist of host antiviral responses. Although NP-NS1 interaction has long been reported, the molecular basis of this protein-protein interaction and its relevance in the virus life cycle remain elusive. Here, we define the NP–NS1 interaction interface and uncover a noncanonical role of NS1 in regulating RNP assembly and viral genome replication. Using integrated biochemical, biophysical, and structural analyses, we demonstrate that the RNA-binding domain (RBD) of NS1 directly engages the NP tail-loop, conferring high affinity for monomeric NP. Functional assays reveal that a cytoplasm-restricted NS1 variant acts as an NP-specific chaperone by stabilizing monomeric NP through sequestration of its intrinsically flexible tail-loop and by scaffolding NP–NP interactions. During influenza virus infection, phosphorylation at the homotypic interface restricts NP in a replication-competent monomeric state. Cytoplasmic NS1 recruits these phosphorylated monomeric NPs into assembling RNPs, thereby facilitating viral genome replication and virus propagation. Collectively, our findings establish NS1 as a selective cytoplasmic chaperone of NP and reveal a novel regulatory axis governing influenza virus RNP biogenesis and replication.

## Introduction

The influenza A virus nucleoprotein (NP) is a major structural component that encapsidates viral genomic RNA in oligomeric form and, together with the heterotrimeric RNA-dependent RNA polymerase (RdRp), forms ribonucleoprotein (RNP) complexes—central machinery driving viral transcription and genome replication (1). The non-structural protein 1 (NS1) is a multifunctional virulence factor and a potent antagonist of host innate immune response, required for optimal virus propagation inside its host (2, 3). Although NS1 has long been known to interact with NP and associate with viral RNPs, the physiological relevance of this interaction and its role in regulating virus replication remain uncharacterised (4, 5).

Post entry and uncoating, antisense viral RNPs (vRNPs) are transported into the host cell nucleus, where they direct viral gene expression (6). The Resident RdRp associated with incoming RNPs initiates transcription using a capped RNA primer, snatched from the host pre-mRNA, producing capped and polyadenylated viral mRNAs (7). In contrast, genome replication is a two-step process that depends upon newly translated RdRp subunits (PB1, PB2 and PA) and NP (8). In the first step, a newly translated trans-activating polymerase forms a dimer with the resident polymerase to form a dimeric replication complex, replicase (9). Within replicase, one RdRp catalyses the synthesis of complementary RNA (cRNA), while the second RdRp promotes the co-replicative encapsidation by recruiting NP to the nascent RNA, thereby generating complementary RNPs (cRNPs) (10, 11). These cRNPs subsequently serve as templates for the production of progeny vRNPs through an analogous mechanism during the second stage of replication (7, 9, 10). Efficient co-replicative assembly of c/vRNPs requires coordinated action of a number of host proteins that aid polymerase dimerization (11), chaperone NP oligomerization (12–15) and post-translationally modify NP (16–18) to regulate their assembly into functional RNP complexes.

Similar to other negative-sense RNA viruses, influenza NP oligomerizes and binds RNA non-specifically (19–21). The atomic structure of different influenza virus NPs (A/H1N1, H3N2, H5N1, B and D) revealed a bilobed architecture, with a head and a body domain connecting to a flexible tail-loop (19, 20, 22–24). In oligomeric NP, this tail-loop from one protomer gets inserted into a groove in the body domain of the neighbouring protomer, forming a key salt-bridge interaction between R416 (tail loop) and E339 (groove) residues (19). A positively charged surface in between the head and body domains accommodates 20-24 nucleotides of RNA (19, 25, 26). These protein-protein and protein-RNA interactions drive NP homo-oligomerization and its assembly into functional RNP complexes (1, 27, 28). However, a downside of these activities is that NP non-specifically binds cellular RNAs to form non-functional aggregates (29, 30). We have shown earlier that reversible phosphorylation of NP at the tail loop–groove interface blocks NP oligomerization, thus help maintaining a monomeric pool of NP required for the RNP assembly process (17, 18). Yet, the precise mechanism by which these monomeric phospho-NP molecules are sequentially recruited on the nascent RNA chain to assemble progeny RNPs remains elusive.

The eighth segment of the influenza virus genome codes for two non-structural proteins, NS1 and NS2, through alternative splicing (31). The NS1 protein is one of the early expressed genes (along with PB1, PB2, PA and NP) and is known to serve as the key factor in antagonizing the host innate immune response, shutting off host gene expression and triggering pathogenesis (3, 32). The protein harbours an N-terminal RNA binding domain, which interacts with various host cellular and viral RNAs, followed by an effector domain that interacts with 30 kilodalton cleavage and polyadenylation factor (CPSF30) to block host mRNA processing, and a C-terminal disordered tail that has been shown to be a critical virulence factor for certain strains of influenza A virus (33–36). Apart from its role in modulating the host, NS1 is known to fine-tune RNP activity and viral RNA synthesis and also regulate vRNP packaging into the virions (37–39). Additionally, NS1 has been shown to associate with the double-stranded RNA loops protruding out of the RNP structures, thereby masking them from the innate immune sensors like RIG-I (40). Very recently, the cryo-EM structure of the influenza virus confirmed the incorporation of NS1 within virion particles, possibly through specific association with the RNP-associated proteins (41). All these observations trace back to the earlier reports suggesting physical association of NS1 with RNPs, possibly through its direct interaction with NP (5). But, the NP-NS1 interaction has never been characterised in molecular detail, thereby keeping the mechanism of NS1-mediated modulation of RNP activity enigmatic.

Here we report a comprehensive biochemical, biophysical, and structural analysis of the NP–NS1 interaction, revealing a non-canonical role of NS1 as a cytoplasmic chaperone of NP. Our data suggests that NS1 specifically interacts with the NP tail-loop domain, restricting its intrinsic flexibility and increasing the stability of monomeric NP structure. In addition, NS1 functions as a scaffold to promote NP tail loop - groove interaction, thereby facilitating NP oligomerization, RNP assembly and viral genome replication. Using an NS1 mutant that predominantly localize into the cytoplasm, we demonstrate that the cytoplasmic NS1 interacts with and stabilises the phosphorylated monomeric NP molecules; subsequently these NS1-NP complexes participate in functional RNP assembly inside the nucleus. Together, our data unravel a critical regulatory role of the NS1 protein in promoting viral genome replication and shed light upon the complex process of dynamic NP oligomerization into a functional RNP complex.

## Results

### NP and NS1 proteins engage in direct interaction in both nuclear and cytoplasmic compartments

To investigate the NS1’s association with NP/ RNP complexes in nfluenza virus infected cells (4, 5), A549 cells were infected with influenza A/H1N1/WSN/1933 virus, and NP/RNP complexes were immunopurified using NP specific antibody (Figure 1A). Viral NP/RNPs copurified the NS1 protein, thereby reconfirming previous reports. Immunofluorescence analysis revealed that both NP and NS1 share similar spatiotemporal distribution with exclusive nuclear accumulation at the early phase of infection (2 h.p.i.) and majorly cytoplasmic localisation at later phases of infection (4 h.p.i. & 6 h.p.i.) (Figure 1B). To check if NP-NS1 colocalization promotes physical interaction between these two proteins at different subcellular spaces, infected cells were subjected to nuclear-cytoplasmic fractionation at 6 h.p.i., followed by immunoprecipitation using NP specific antibody. As evidenced, NP-NS1 complexes were immunopurified both from nuclear and cytoplasmic fractions (Figure 1C). This is in contrast to the previous report where NS1-RNP complexes were isolated using PB2 (RdRp) specific antibody, exclusively from the nuclear chromatin factions (5). This is suggestive of the fact that NS1 may interact with free NP in the cytoplasm while associating with NP/RNPs inside the nucleus. To validate this data, we utilised a mutant NS1, NS1-R148A/E152A/E153A (NS1-AAA), which was previously reported to preferably localize in the cytoplasm of the infected cells (42, 43). As shown, NS1-AAA majorly accumulated in the cytoplasm while the WT NS1 preferentially localized in the nucleus when expressed in cells through transient transfection (Figure 1D & 1E). The NS1-AAA immunoprecipitated NP comparable to the WT NS1, thereby validating a cytoplasmic one-to-one interaction between NP and NS1, independent of other RNP components (Figure 1F). Notably, this cytoplasmic variant of NS1 can also localize into the nucleus upon co-expression with NP (Figure S1A), suggesting NP-NS1-AAA complexes, although formed in the cytoplasm, can subsequently translocate to the nucleus.

**Figure 1:**
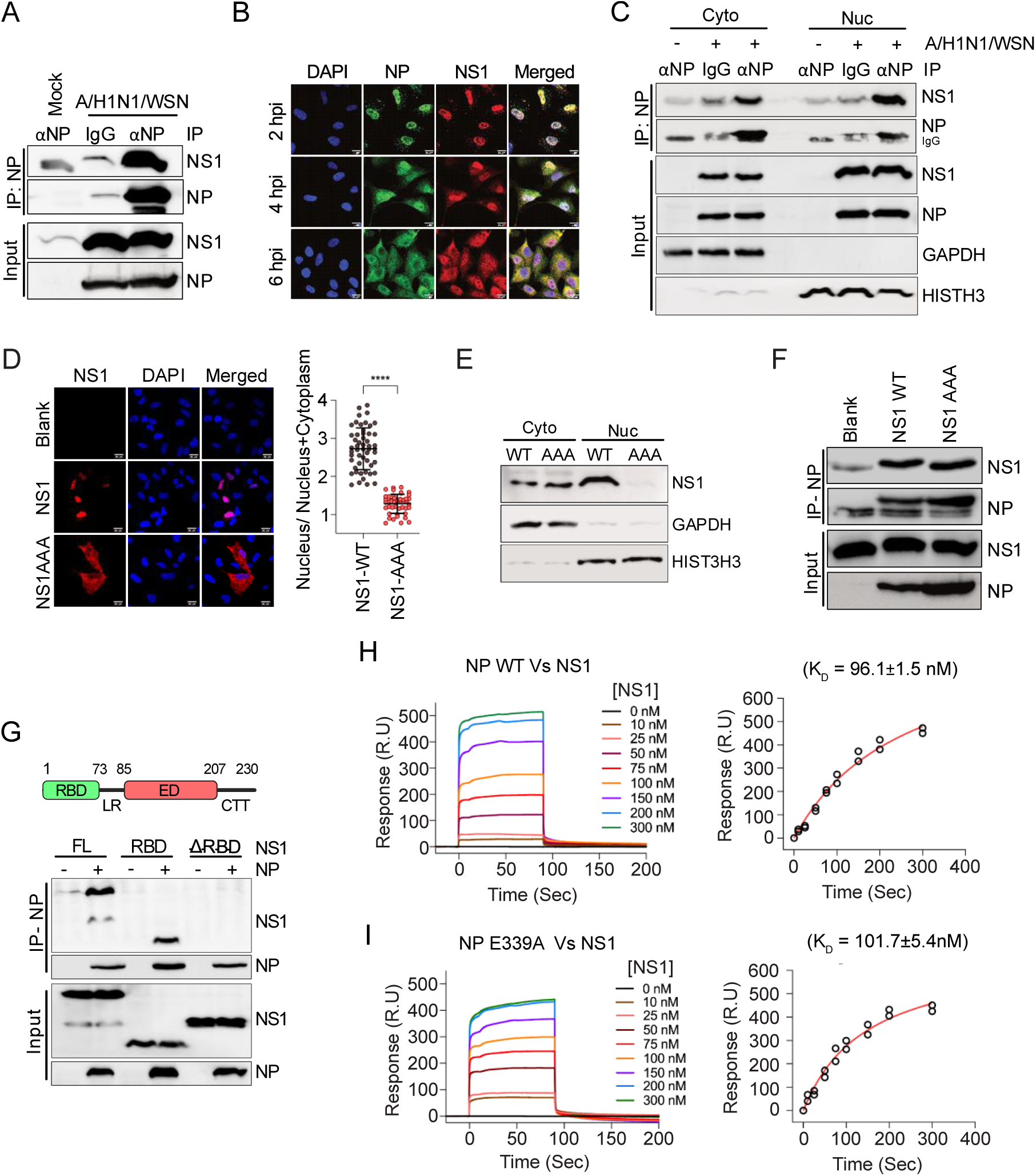
NP and NS1 proteins engage in direct interaction in both nuclear and cytoplasmic compartments. (A) A549 cells infected with IAV/WSN/33 (MOI 0.1) were harvested 24 h post-infection, and NP was immunoprecipitated, followed by immunoblotting for NP and NS1. (B) Infected A549 cells (MOI 5) were fixed and stained for NP and NS1 and imaged through a confocal microscope. (C) Infected A549 cells (MOI 2) were harvested 6 h.p.i., followed by nuclear-cytoplasmic fractionation and immunoprecipitation using NP-specific antibody. (D-E) FLAG-tagged NS1 WT or NS1-AAA was expressed in A549 cells, and subcellular localization was analyzed by confocal microscopy (D) and nuclear/cytoplasmic fractionation (E). For D, nuclear-cytoplasmic distribution of NS1 in transfected cells (n=50) was quantified in ImageJ. (F-G) NP-V5 was co-expressed with FLAG-tagged NS1 WT or NS1-AAA (F) or with WT or truncated variants of FLAG-tagged NS1 (G), followed by IP using V5 antibody and blotting with V5 (NP) and FLAG (NS1) antibodies. (H-I) SPR analysis of NP WT (H) or NP E339A (I) immobilized on CM5 chips with varying concentrations of NS1 FL to determine binding affinity (K_D)_.

To characterize the NP-NS1 interaction in molecular detail, we generated truncated variants of NS1 lacking either the N-terminal RBD or the C-Terminal effector and tail domains. The RNA-binding domain of NS1 appeared to be self-sufficient for interaction with NP as evidenced through co-immunoprecipitation assay (Figure 1G). Subsequently, we expressed and purified recombinant NP, full-length (FL) and the RBD domain of NS1. During size exclusion chromatography, NP eluted as a combination of oligomeric and monomeric forms (Figure S1B) (17). NS1-FL was distributed as a predominant tetrameric population, while NS1-RBD was eluted as a dimer (figure S1C & D) (33, 44). To exclude the possibility of NP oligomerization interfering with its ability to interact with NS1, we also purified the obligatory monomeric NP E339A mutant (Figure S1B) (19). All the protein preparations were devoid of non-specifically bound cellular RNA, as confirmed by the OD-260/280 ratio ranging between 0.5-0.6. His-tag coelution assay showed that NS1-RBD co-elutes with his-tagged NP (figure S1 E), thereby confirming that NP and NS1 proteins are bonafide interacting partners. Subsequently, we employed surface plasmon resonance (SPR) to estimate the affinity of the NP-NS1 interaction. NS1-FL or NS1-RBD were injected as analytes over the immobilised NP-WT or NP-E339A. Increasing concentration of the analyte resulted in increasing binding response, accounting for a dissociation constant of 96 nanomoles for the interaction between WT NP and NS1-FL (Figure 1H); monomeric NP E339A showed binding affinity (101 nM) comparable to WT NP (Figure 1I). The NS1 RBD showed lower binding affinity towards NP WT and E339A (71 µM and 89 µM respectively) compared to the full-length NS1(Figure S1F & G). This data suggested that although the NS1 RBD domain serves as the primary site of interaction with NP, other domains of NS1 also synergize to establish a stronger interaction between these two proteins. Hence, all subsequent experiments were performed exclusively with full-length NS1 protein.

### NP-NS1 participates in a bimodal interaction

To delineate the NP domains involved in interaction with NS1, we analyzed a panel of NP variants differing in oligomeric state and tail loop accessibility. The panel included the monomeric NP-E339A mutant, the previously reported NP-S486A mutant that exists exclusively as oligomers, and an NP variant lacking the entire tail loop (NPΔTL), which also remains monomeric (45). Size-exclusion chromatography confirmed the respective oligomerization state of these NP variants (Figure 2A). All proteins were assessed for NS1 binding by SPR. The monomeric NP-E339A bound NS1 with the highest affinity (K_D =_ 0.12 μM) (Figure 2B, S2A). In contrast, NPΔTL exhibited a ∼25-fold reduction in binding affinity (K_D =_ 3.01 μM) (Figure 2C, S2B), indicating the critical role of the NP tail loop in interaction with NS1. Notably, NS1 showed moderate binding to oligomeric NP-S486A (K_D =_ 0.9 μM; ∼7.5-fold weaker than NP-E339A) (Figure 2D, S2C), in which the tail loop remains buried within the groove of the neighbouring protomer and hence should remain inaccessible to NS1 (Figure 2D, upper panel). Collectively, these results suggest that NS1 engages monomeric NP through a tail loop–dependent interaction, while for oligomeric NP it interacts with an alternative interaction interface devoid of the tail loop domain.

**Figure 2:**
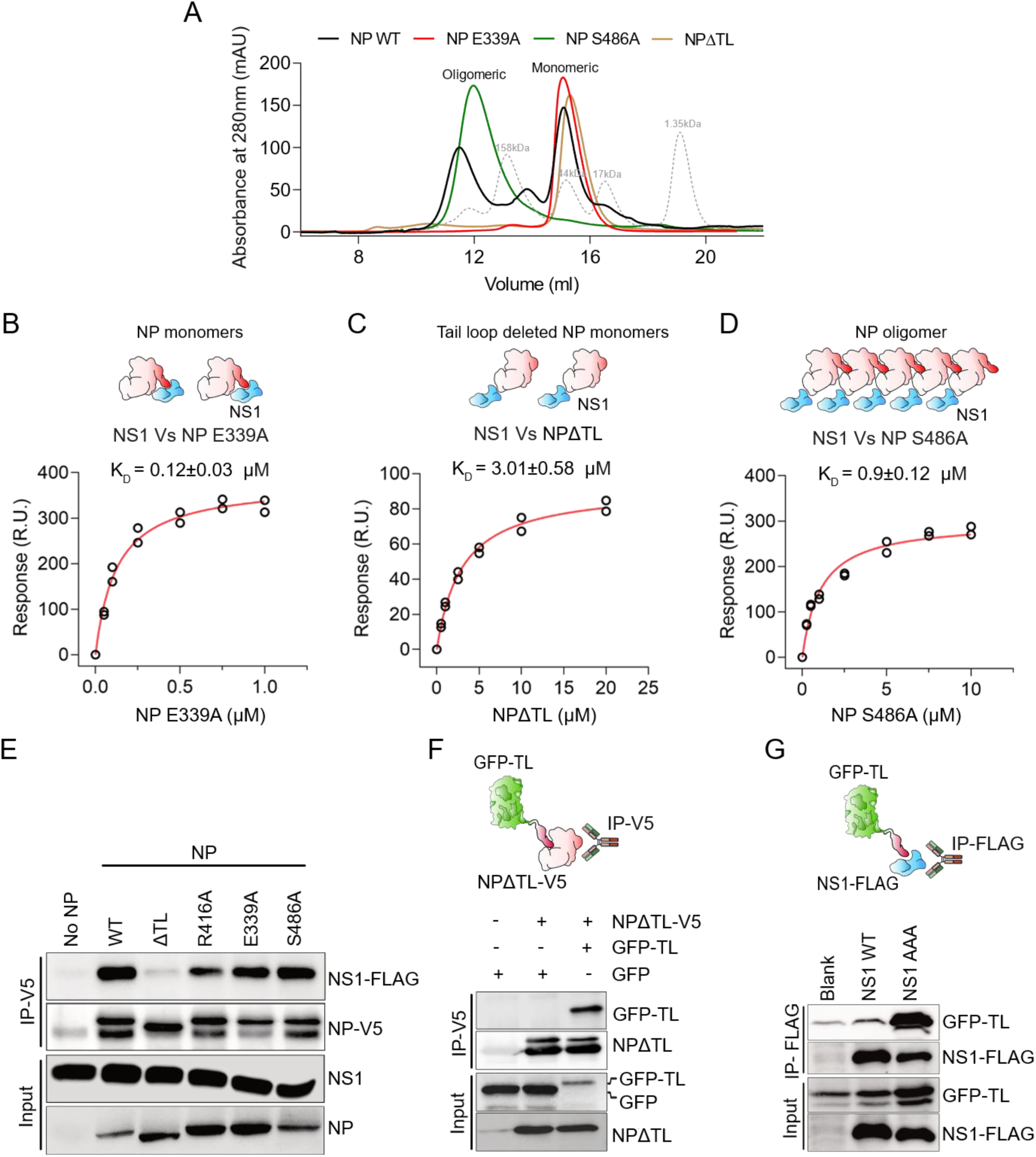
Bimodal nature of NP-NS1 interaction. (A) Size exclusion chromatography profiles of purified NP WT and mutants (E339A, S486A, and ΔTL) resolved on ENrich SEC650. (B-D) SPR analysis of NS1 immobilized on CM5 chips with NP mutants E339A (B), ΔTL (C), and S486A (D) as analytes; affinity constants (K_D)_ were determined by fitting in Biacore instrument software. Schematic representation of different NP (red-white)-NS1 (blue) interactions. (E) V5-tagged NPs WT or mutants (ΔTL, R416A, E339A, S486A) were co-expressed with FLAG-NS1 in HEK293T cells and analyzed by co-IP. (F) V5-tagged NPΔTL and GFP-TL were co-expressed in HEK293 cells, and protein-protein interactions were analyzed by IP using V5 antibody followed by blotting with GFP and V5 antibodies. (G) GFP-TL was co-expressed with FLAG-tagged NS1 WT or NS1-AAA in HEK293T cells and analyzed by co-IP using anti-FLAG beads followed by blotting using GFP antibody.

Co-immunoprecipitation assay was performed to validate the SPR findings in the cellular context. V5 epitope-tagged monomeric NP-E339A and oligomeric NP-S486A co-precipitated FLAG-tagged NS1 at levels comparable to wild-type NP (Figure 2E). Interestingly, mutation in tail loop residue R416 to alanine (NP-R416A) showed markedly reduced NS1 co-precipitation, whereas complete deletion of the tail loop severely affected the interaction (Figure 2E). Treatment of the cell lysate with RNaseA before immunoprecipitation led to similar results thereby excluding any potential role of RNA in mediating NP-NS1 interaction (Figure S2D). Together these results corroborate the SPR data and further validate the critical role of the NP tail loop in mediating monomeric NP–NS1 interaction. To assess if the NP tail loop directly participate in interaction with NS1, we employed a chimeric GFP fused to the NP tail loop at its C-terminus (GFP-TL) (17). When co-expressed in cells, GFP-TL co-precipitated with V5-tagged NP lacking the tail loop domain-NPΔTL (Figure 2F), thus confirming that the isolated tail loop remains functionally competent to participate in the tail loop–groove interaction. GFP-TL co-precipitated with FLAG-tagged NS1 (Figure S2E), demonstrating that the NP tail loop alone is sufficient to mediate a direct interaction with NS1. Notably, the cytoplasmic NS1-AAA showed around a 5-fold enhanced co-precipitation of GFP-TL (Figure 2G), hence establishing the cytoplasmic NS1 as a superior interacting partner for the NP tail loop domain. This might also be due to the colocalization of both the chimeric GFP-TL and the NS1-AAA preferentially in the cytoplasm while NS1 majorly localizes in the nucleus. Nonetheless, these data indicate that NS1 can stably interact with monomeric NP through its tail loop domain, and possibly the interaction occurs in the cytoplasm.

### NS1 acts as a cytoplasmic chaperone to stabilise monomeric NP

Post translation, NP gets phosphorylated at its homotypic interface that blocks its oligomerization and maintains the protein in monomeric form (16, 17), which harbors a free and highly flexible tail loop domain (46), adopting different structural conformations (Figure S3A). To test if interaction with NS1 may influence the structural stability of the tail loop and the monomeric NP as a whole, we performed molecular dynamic simulation. Structural dynamics of a single protomer of the trimeric NP were analysed either alone or in complex with NS1. Our data showed higher RMSD and Rg values for the free monomeric NP promoter compared to when it remained in complex with NS1 (Figure 3A & B, Table 1), suggesting compaction of the NP structure upon complex formation with NS1. A distinct effect was observed for the NP tail loop (residues 402–428). In the monomeric state, this region displayed high mobility, as evidenced by higher RMSD, indicative of a disordered and transiently collapsed ensemble (Figure 3D, Table.1). This is in corroboration with the crystal structure of monomeric NP (NP R416A) where the tail loop, instead of binding to the groove of the neighboring protomer, collapses onto its own RNA-binding surface (47). Upon complexation with NS1, the tail loop became substantially more stable as evidenced from the decrease in the standard deviation of Rg from 0.09 to 0.03 nm (Figure 3E, Table 1). This further supports conversion of this region from a fluctuating disordered segment into a structurally restrained recognition element. This is also supported by the residue-wise RMSF values of NP which elucidated higher degrees of molecular motion of the tail loop domain in monomeric NP, which was completely lost in the NP-NS1 complex (Figure 3C). Additionally, RDF analysis of water molecules around the NP tail loop showed a pronounced reduction in solvent density in the complex (Figure 3F). These results indicate that the tail loop, which is highly mobile and solvent-exposed in the monomeric state, becomes more ordered, slightly extended, and less hydrated upon NS1 binding.

**Figure 3:**
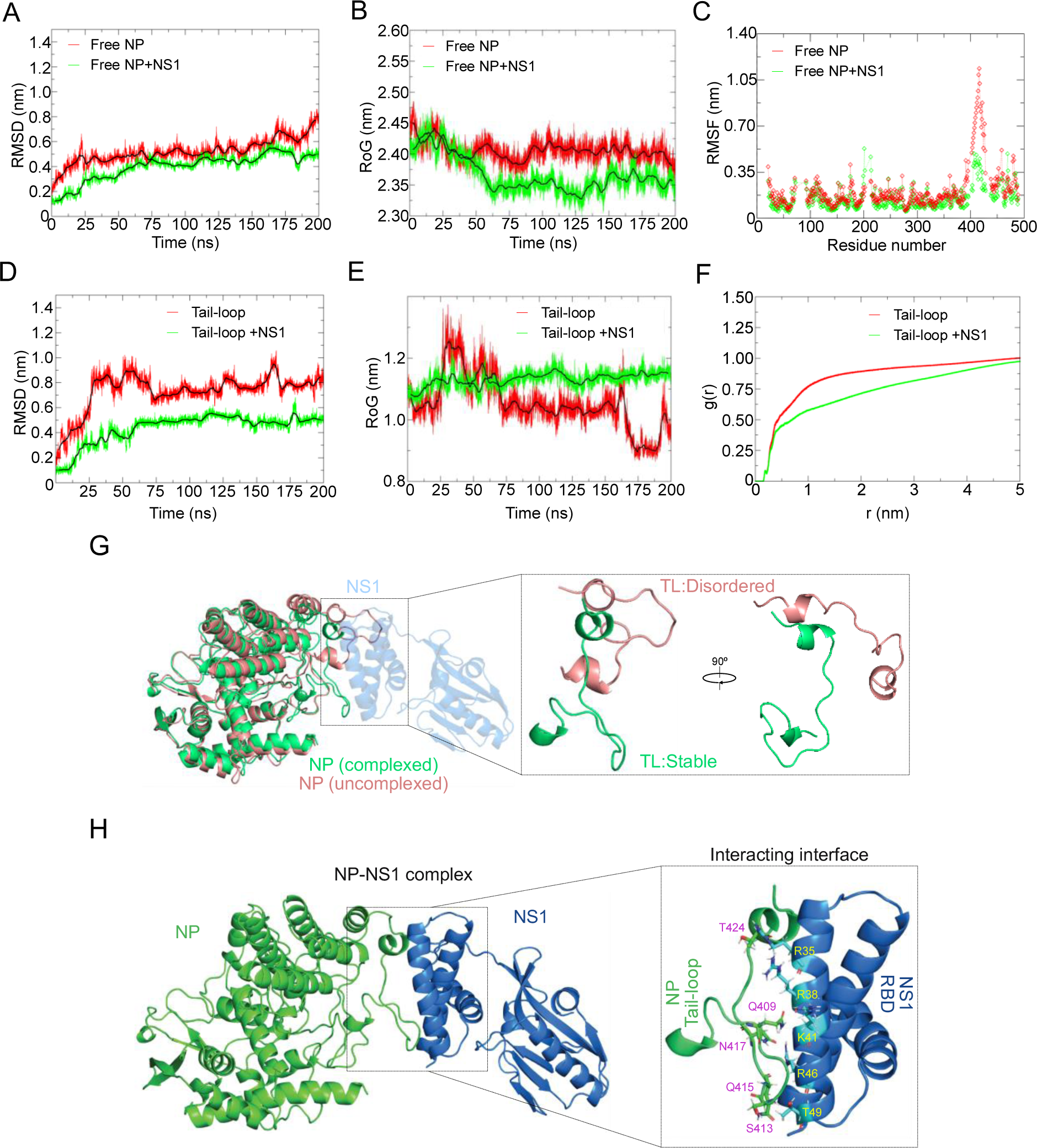
NS1 stabilizes NP monomer structure by reducing flexibility of the tail loop domain. (A-F) MD simulations comparing free and NS1-bound NP monomer, showing backbone RMSD (A), Rg (B) and RMSF (C) of the whole NP structure (individual chain of trimeric NP, 2iqh) or the RMSD (D), Rg (E), and water radial distribution function (RDF) (F) of the tail loop domain. (G) Structural superposition of representative conformations of free monomeric NP (salmon) or the NS1 (blue)-bound NP (green). (H) Global minimum FEL of the NP-NS1 complex highlighting major interaction interfaces. NP (green) and NS1 (blue) residues are marked in violet and yellow, respectively.

**Table 1:** Comparison of the RMSD and Rg values of NP and NP tail-loop domain both in free and bound state.

| Parameter | Free NP (Mean ± SD) | NP-NS1 Complex (Mean ± SD) |
| --- | --- | --- |
| Backbone RMSD (nm) | 0.52 ± 0.10 | 0.40 ± 0.11 |
| Radius of Gyration (Rg) (nm) | 2.41 ± 0.02 | 2.37 ± 0.03 |
| Tail-loop RMSD (nm) | 0.73 ± 0.15 | 0.44 ± 0.12 |
| Tail-loop Rg (nm) | 1.05 ± 0.09 | 1.13 ± 0.03 |

We further explored the interaction interface between NP and NS1, retrieving from the structure lying at the energy minima of the free energy landscape (FEL). The results revealed that the monomeric NP-NS1 interaction was driven by a series of high-occupancy hydrogen bonds mediated by NS1—Arg35, Arg38, and Arg46 residues with Glu412, Ser413, Gln415, Asn417 and Thr424 residues of NP. This arginine-rich anchoring is further stabilized by a contiguous hydrophobic patch of NS1 (Arg41, Leu42 and Pro43), which packs against a nonpolar cluster on NP (Ile414, Met425, and Val421) (Figure 3H, S3B). Notably, structural orientation of the tail-loop domain revealed that complex formation with NS1 preserves the tail-loop domain in its native functional conformation that is favorable for efficient NP oligomerization, whereas in the unbound state it appears distorted and deviates from its native structural arrangement (Figure 3G). All these in-silico results strongly suggested that complex formation with NS1 can stabilize the monomeric NP structure by reducing the flexibility of the tail loop domain.

To validate the in-silico MD data, we took two different approaches. First, we tried validating the NP-NS1 interaction interface as elucidated by the in-silico analysis. The critical NP-interacting residues of NS1, R38 and K41, were mutated to alanine. As observed, the mutant NS1 failed to interact with NP in co-immunoprecipitation analysis (Figure 4A). Subsequently, recombinant NS1 WT or the R38A/K41A mutants were expressed, purified (Figure S4A) and tested for their comparative binding affinity towards monomeric NP E339A or oligomeric NP S486A using SPR. The NS1 mutant showed severely compromised binding affinity towards the monomeric NP (5-10-fold less than the WT) (Figure 4B, S4B & C, D & E), while binding to the oligomeric NP remained unaffected (Figure 4C, S4F & G). This data confirmed that NS1 utilizes R38 and K41 residues for selective interaction with the tail loop domain of monomeric NP, while it employs a distinct interface (excluding the R38 and K41 residues) for binding to oligomeric NP.

**Figure 4:**
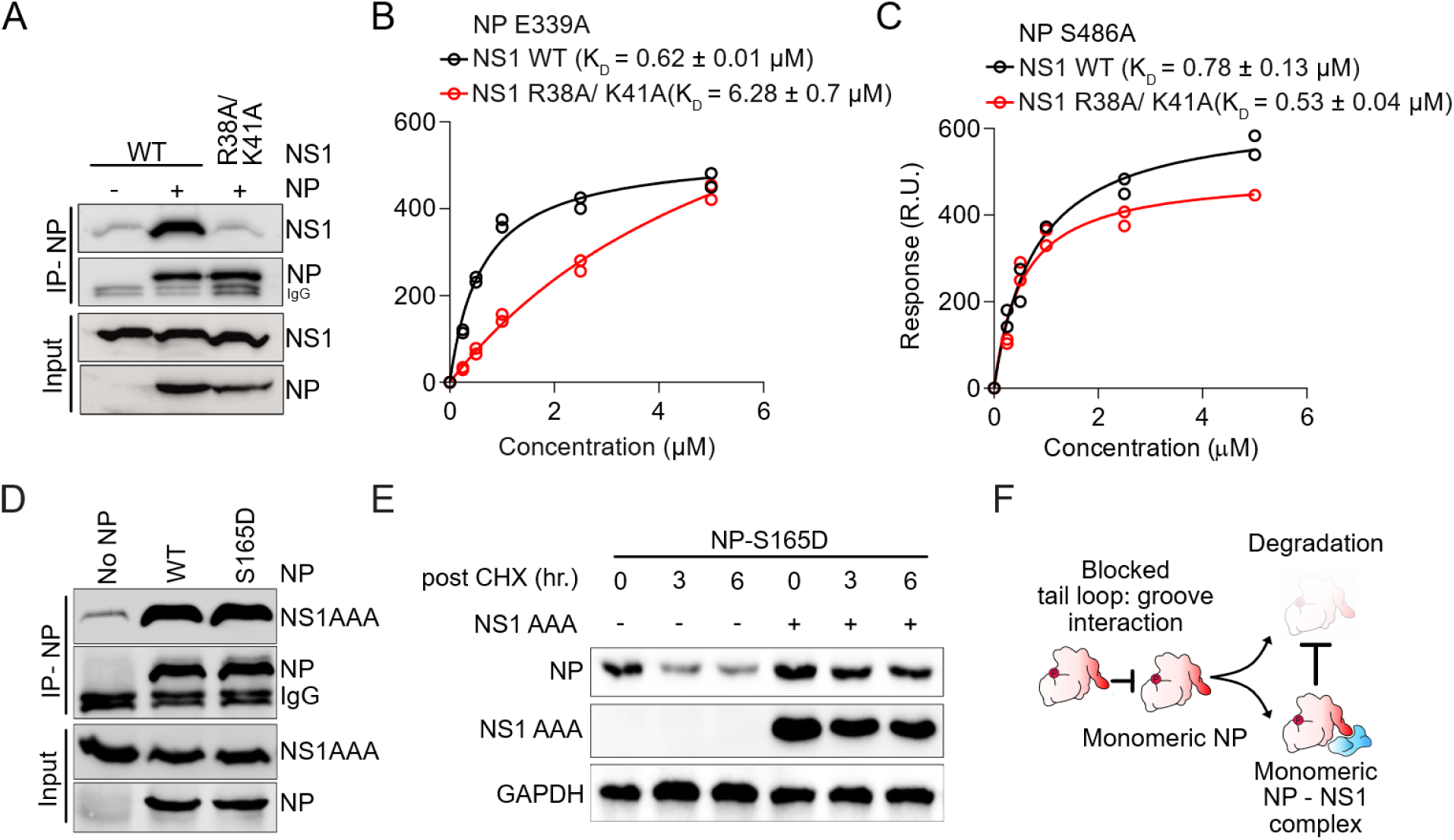
NS1 acts as a cytoplasmic chaperone to stabilize monomeric NP. (A) NP-V5 was co-expressed with FLAG-tagged NS1-WT or NS1-R38A/K41A in HEK293T cells. Co-IP was performed using anti-V5 antibody. (B-C) SPR analysis of NP E339A (B) and NP S486A (C) with NS1 WT or NS1 R38A/K41A to assess binding affinity (K_D)_. (D) V5-tagged NP WT or NP S165D was co-expressed with NS1-AAA in HEK293T cells and analyzed by co-IP using V5 antibody. (E) NP S165D was co-expressed with NS1-AAA in HEK293T, followed by cycloheximide chase and immunoblotting with NP and NS1 antibodies. GAPDH serves as an internal control. (F) Schematic model showing the fate of phosphorylated NP monomers in the absence or presence of cytoplasmic NS1.

Next, we assessed the in-cell stability of monomeric NP in the presence or absence of cytoplasmic NS1-AAA. Phosphorylation of NP within the tail loop binding groove, at Ser165, sterically blocks the tail loop-groove interaction and keeps the protein in monomeric form (16, 17). Hence, the NP S165D mutant remains in monomeric form in vitro and in cells (Figure S5A & (22) respectively). We assessed if NS1-AAA can interact with and hence influence the stability of this monomeric NP S165D. NS1-AAA interacted with NP-S165D comparable to the WT NP in co-immunoprecipitation assay (Figure 4D). When co-expressed, NS1-AAA greatly enhanced the in-cell stability of the NP S165D, as evidenced by the cycloheximide chase experiment, thus confirming its chaperone activity towards the monomeric phospho-NP molecules (Figure 4E & F). These experiments, along with the in-silico data, together establish cytoplasmic NS1 can act as an NP specific chaperone through its direct interaction with the free tail loop domain, reducing its inherent flexibility and stabilizing the monomeric NP structure.

### NS1 supported NP oligomerization model

Given that NS1 interacts with the NP tail loop domain that governs homotypic interaction of NP, we examine whether NS1 modulates NP oligomerization, either in a positive or negative way. Binding free energies for the NP-NS1 heterodimer and NP-NP homodimer were calculated using the MM/PBSA method. The MM/PBSA results indicate that both complexes are energetically stable, but the NP-NP homodimer (ΔG_bind =_ −88.63 ± 13.78 kcal/mol) possesses a binding affinity much higher than that of the NP-NS1 heterodimer (ΔG_bind =_ −4.48 ± 11.12 kcal/mol) (Table 2). This suggests that NP self-association is thermodynamically preferred over the formation of the NP-NS1 complex. Thus, we speculated that NS1 interacts with tail-loop domain of NP monomers, stabilizing them in a conformation favorable to oligomerization. This in turn facilitates NP oligomerisation, as the local concentration of NP monomers increases (Figure 5A).

**Figure 5:**
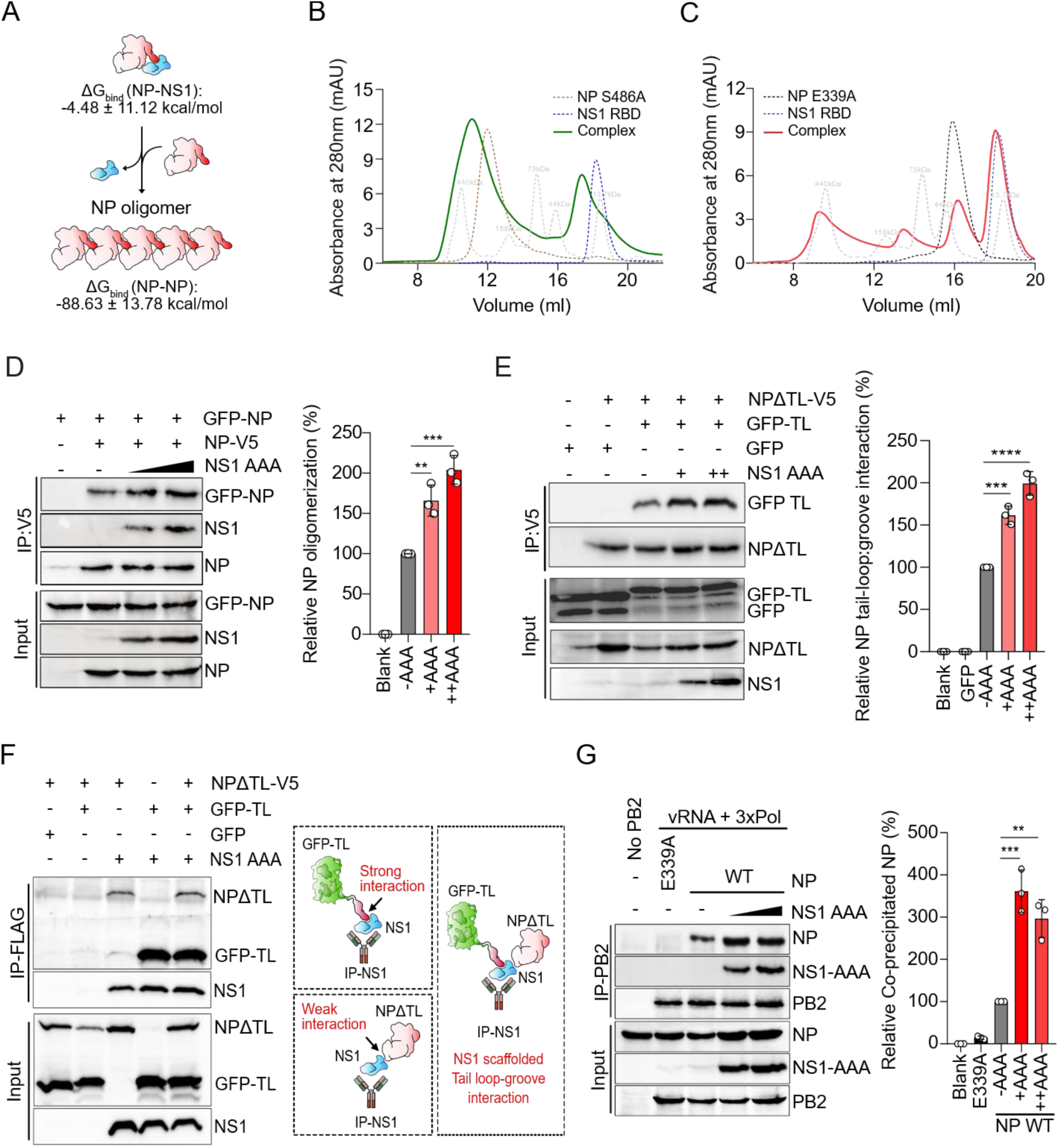
NS1 supported NP oligomerization model. (A) Model illustrating comparative interaction energies of NP-NS1 and NP-NP interactions. (B-C) SEC analysis of complexes formed between NS1 RBD and NP S486A (B) or NP E339A (C) on a Superdex 200-10/300 GL column. (D) NP-V5 and NP-GFP were co-expressed in HEK293T cells in the presence of increasing concentrations of FLAG-tagged NS1-AAA. Homotypic interaction was monitored by IP using V5 antibody and blotting using GFP and V5 antibodies. (E) NPΔTL-V5 and GFP-TL were co-expressed with or without FLAG-NS1-AAA. Tail loop-groove interaction was monitored by IP using V5 antibody and blotting using GFP and V5 antibodies. (F) NPΔTL-V5 and GFP-TL were co-expressed with NS1-AAA-FLAG either individually or together, and protein-protein interactions were analyzed through IP using FLAG antibody and blotting with V5 and GFP antibodies. Cartoon illustrating the co-immunoprecipitation results. (G) Viral RNPs were reconstituted by co-expressing viral RNA, NP and RdRp subunits in the absence and presence of increasing concentrations of NS1-AAA. The extent of RNP assembly was measured by IP using PB2 antibody and blotting with NP antibody. For D, E and G, band intensities were quantified in Image Lab software (Bio-Rad) and plotted as a relative percentage of the no-NS1 control. Statistical significance was analyzed by unpaired two-tailed equal variance. Student’s t-test between the individual sets with P values: ns>0.05, *P<0.05,* *P<0.01, ***P<0.001, and ****P<0.0001.

**Table 2:** Free energy of binding for the NP-NS1 heterodimer and the NP-NP homodimer.

| Energy Component | NP-NP<br>(Average ± SD)<br>(kcal/mol) | NP-NS1<br>(Average ± SD)<br>(kcal/mol) |
| --- | --- | --- |
| Van der Waals energy (ΔVDWAALS) | -211.78 ± 8.14 | -80.03 ± 6.93 |
| Electrostatic energy (ΔE<sub>EL</sub>) | 840.64 ± 71.90 | -549.23 ± 42.49 |
| Polar solvation energy (ΔE<sub>PB</sub>) | -790.70 ± 68.15 | 553.71 ± 39.37 |
| Non-polar solvation energy (ΔE<sub>NPOLAR</sub>) | -21.92 ± 1.02 | -10.54 ± 0.53 |
| Gas-phase energy (ΔG<sub>GAS</sub>) | 628.86 ± 70.01 | -629.26 ± 41.20 |
| Solvation free energy (ΔG<sub>SOLV</sub>) | -812.62 ± 67.82 | 543.18 ± 39.28 |
| Total binding free energy (ΔTOTAL) | -183.76 ± 13.78 | -86.08 ± 11.12 |
| Entropy-corrected binding free energy (ΔG<sub>bind</sub>) | -88.63 ± 13.78 | -4.48 ± 11.12 |

To validate this, we assessed potential alteration of the oligomerization state of monomeric NP-E339A and oligomeric NP-S486A upon complex formation with NS1, as analyzed by size-exclusion chromatography. In the absence of NS1, NP-S486A displayed a characteristic oligomeric elution profile, which shifted modestly toward higher molecular-weight fractions upon NS1 binding (Figure 5B). In contrast, monomeric NP-E339A eluted as a heterogeneous population of higher-order species when complexed with NS1 (Figure 5C). These results indicate that NS1 does not inhibit, rather promotes oligomerization of monomeric NP. To validate this further, NP self-association was monitored in cells in the presence and absence of NS1-AAA. As observed, V5-tagged NP (NP-V5) can be efficiently coprecipitated with N-terminal GFP-fused NP (GFP-NP), confirming homotypic interaction. Notably, increasing amounts of NS1-AAA resulted in a dose-dependent enhancement of GFP-NP coprecipitation, establishing the oligomerization-promoting effect of NS1 (Figure 5D). Interestingly, NS1-AAA was also detected in the immunoprecipitated complex, suggesting that NS1 remains associated with NP homo-oligomers. To pinpoint the molecular mechanism underlying NS1’s NP oligomerization-promoting activity, we reconstituted the tail loop-groove interaction by co-expressing the GFP-TL and NPΔTL in the absence or presence of NS1-AAA. Cytoplasmic NS1 enhanced GFP-TL coprecipitation with NPΔTL in a concentration-dependent manner (Figure 5E). Additionally, NS1 was also able to coprecipitate GFP-TL and NPΔTL together in a separate co-immunoprecipitation assay, thus supporting a model where NS1 bridges the tail-loop–groove interaction and acts as a molecular scaffold to facilitate homotypic interaction between individual NP molecules (Figure 5F). Notably, GFP-TL showed greater co-precipitation than NPΔTL, corroborating the SPR data and indicating that NS1 shows stronger affinity towards the NP tail loop domain.

Finally, we assessed if cytoplasmic NS1 (NS1-AAA) could also facilitate assemblies of viral RNPs inside the nucleus. Viral RNPs were reconstituted in cells by expressing the RdRp subunits (PB2-HA, PB1 and PA), WT or mutant NP, and a vRNA template. The efficiency of RNP formation was monitored by immunoprecipitating the RdRp via PB2-HA and detecting co-precipitated NP as a part of RNP. Large extent of WT NP was co-purified with RdRp, indicating efficient RNP formation. The presence of NS1-AAA significantly boosted RNP assembly, thereby indicating that NS1 not only promotes NP oligomerization but also drives these NP oligomers towards functional RNP assemblies (Figure 5G). Importantly, NS1 also co-precipitated with RNP complexes, suggesting that NS1, while scaffolding the RNP assembly process also gets physically associated with the same.

### Cytoplasmic NS1 promotes viral genome replication in NP dependent manner

NP’s assembly into functional RNP complexes is the key for optimal synthesis of viral RNA. Hence NS1 or its cytoplasmic variant was tested for its ability to modulate viral RNA synthesis. Influenza A virus reporter RNPs were reconstituted in HEK293T cells as discussed previously (48) in the presence or absence of NS1. Increasing NS1 expression resulted in a dose-dependent enhancement of RNP activity for both HA-and NA-segment–based reporter RNAs, indicating that NS1 positively regulates viral RNA synthesis independent of genomic segment identity (Figure 6A & B, S6A & B). Notably, the NS1 RNA-binding domain (RBD), which mediates interaction with NP, was sufficient to stimulate RNP activity, whereas the effector domain (ΔRBD) was inadequate to do so (Figure 6C, S6C). Consistently, the cytoplasmic NS1-AAA mutant enhanced RNA synthesis to a level comparable to wild-type NS1 (Figure 6D, S6D).

**Figure 6:**
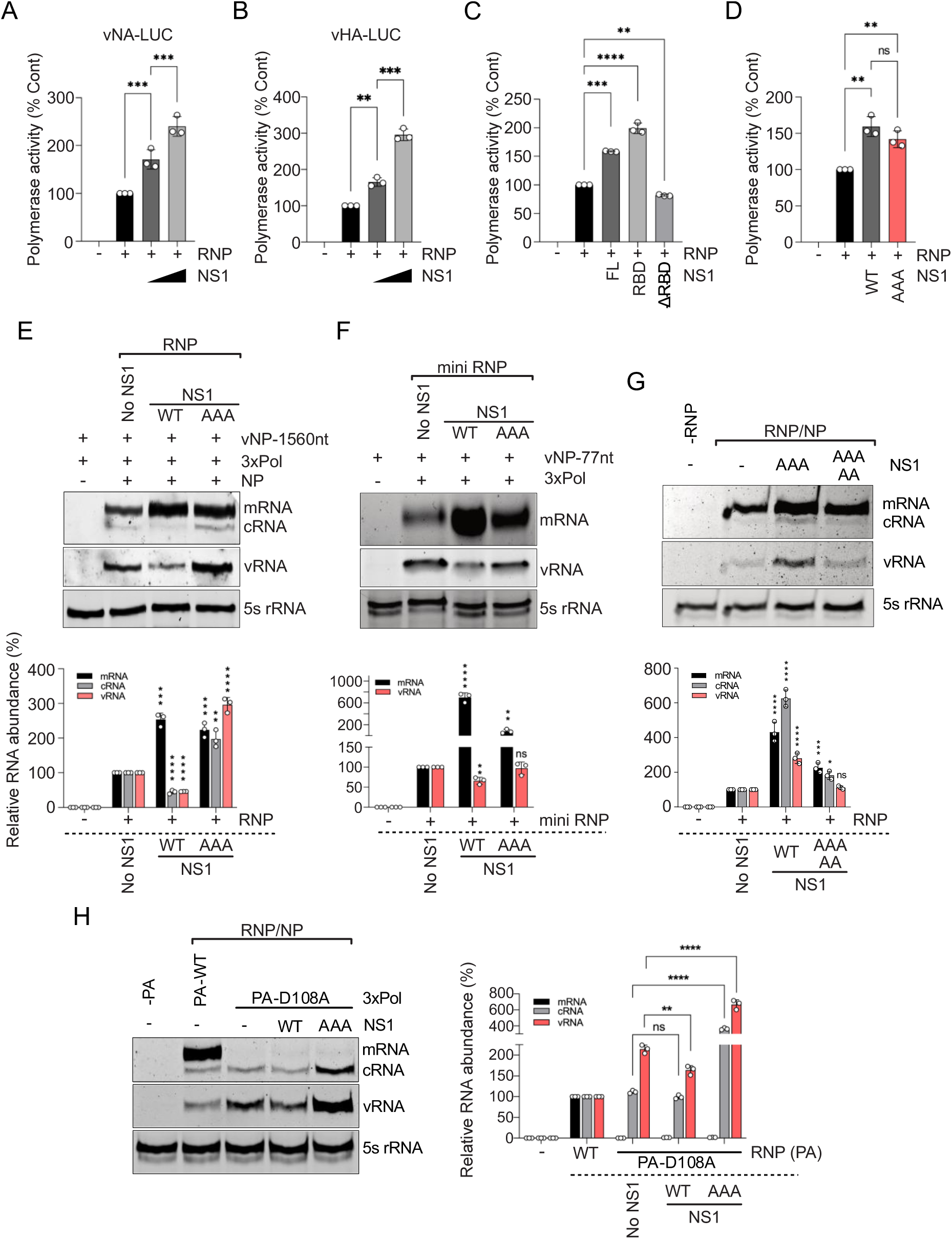
Cytoplasmic NS1 promotes viral genome replication in NP dependent manner. (A-D) Influenza virus (IAV/WSN/33) reporter RNPs were reconstituted in HEK293T cells by co-expressing PB2, PB1, PA and NP proteins and NA (A) or HA (B) segment-based reporter RNA templates in the absence or presence of increasing concentrations of NS1 or full-length or truncated variants of NS1 (NS1 FL or NS1 RBD or NS1 ΔRBD) (C) or NS1-WT or NS1-AAA (D) Reporter activity is plotted as a relative percentage of the no NS1 control. (E-F) Primer extension analysis: full-length RNPs (E) or mini-RNPs devoid of NP (using the NP77 template) (F) were reconstituted in HEK293T cells in the absence or in the presence of NS1 WT or NS1-AAA. Total RNA was isolated, followed by primer extension using mRNA/cRNA and vRNA-specific primers (G) Effect of NS1-AAA 38A/41A on the viral RNA synthesis in the context of reconstituted RNPs observed through primer extension analysis. (H) Effect of NS1-AAA on the viral RNA synthesis in the context of RNPs reconstituted with transcription-defective RdRp (PA-D108A). The band intensity of vRNA, cRNA & mRNA was quantified in Image Lab software. Viral RNAs were normalized to 5s rRNA and plotted as a relative percentage of the no-NS1 control. Each experiment is performed in biological triplicate. Statistical significance was analyzed by two-way ANOVA between the individual sets with P values: ns>0.05, *P<0.05,* *P<0.01, ***P<0.001, and ****P<0.0001.

Primer-extension analysis was performed to investigate which specific step(s) of viral RNA synthesis can be promoted by NS1. Viral RNPs were reconstituted with the NP RNA segment harboring a premature stop codon in the ORF, along with NP, PB1, PB2 and PA proteins, in absence or in presence of WT NS1 or NS1-AAA. Interestingly, NS1 WT and NS1-AAA showed distinct effects upon viral RNA synthesis. WT NS1 boosted mRNA synthesis more than 2.5-fold compared to the control (no NS1) but reduced vRNA and cRNA synthesis to a significant extent (Figure 6E). This is in corroboration with NS1’s transcription-promoting effect as reported earlier (38). In contrast, NS1-AAA upregulated all three RNA synthesis to a great extent, with a maximum increase of vRNA synthesis up to 3-fold compared to the control set (Figure 6E). This data elucidated, for the first time, that NS1, depending on its subcellular localization, can differentially influence viral RNP’s gene transcription and genome replication activities.

To investigate which one of these NS1-mediated effects might depend upon its interaction with NP, we reconstituted NP-free RNPs using the mini-NP77 RNA template that harbors the 3’ and 5’ UTRs of the viral genomic RNA but is devoid of the coding region (49). We and others have shown that RdRp can execute both transcription and replication using this mini-RNA template in an NP-independent manner (48, 49). As evidenced, both NS1 and NS1-AAA promoted transcription, but the latter failed to promote genomic RNA replication in the context of the mini-RNA template. This suggests that the cytoplasmic NS1 can boost viral genome replication in NP dependent manner (Figure 6F). To further validate this, RNPs with full-length NP segments were reconstituted in the presence of NS1-AAA harboring the R38A/K41A mutations, which makes the protein defective in interaction with monomeric NP (Figure 6SE). Primer-extension analysis showed that this NS1-AAA-R38A/K41A mutant was unable to boost viral genome replication, thereby establishing that monomeric NP-NS1 interaction is essential for replication-promoting activity of NS1-AAA (Figure 6G). Finally, we employed a transcription-defective polymerase harboring D108A mutation in the PA endonuclease active site (50). PA-D108A polymerase showed no mRNA synthesis but supported cRNA and vRNA synthesis comparable to the WT polymerase, thereby providing a unique system to monitor viral genome replication in isolation. Overexpression of NS1 resulted in a minor decrease in vRNA synthesis, while overexpression of NS1-AAA greatly elevated both cRNA and vRNA synthesis, thus confirming that NS1-AAA can promote genomic RNA replication independent of transcription (Figure 6H & S6F). Together these data elucidate a dual regulatory effect of NS1 upon viral RNA synthesis; while NS1 in general can promote viral gene transcription, possibly by directly boosting RdRp activity, cytoplasmic NS1 can specifically stimulate genome/antigenome RNA replication, which relies upon NP-mediated RNP assembly process and is rooted in monomeric NP-NS1 interaction.

### Higher cytoplasmic abundance of NS1 promotes v/c-RNA synthesis during influenza virus infection

Next, we intend to validate that higher cytoplasmic abundance of NS1 favors elevated levels of v/c-RNA synthesis during influenza virus infection. Cells exogenously expressing NP, RdRp subunits and either NS1 or NS1-AAA were infected with A/H1N1/WSN virus, followed by cycloheximide treatment. At 6 hours post infection, cells were harvested, followed by RNA isolation and primer extension to evaluate the expression of viral RNA transcripts. While incoming viral RNPs can only execute primary transcription (mRNA) in the presence of cycloheximide, exogenously expressed NP and RdRp proteins supported both transcription (mRNA) as well as replication (cRNA & vRNA). Co-expression of WT NS1 along with the RNP component proteins boosted mRNA levels to a significant extent, thereby confirming the transcription-promoting effect of NS1. Notably, WT NS1 can also elevate both cRNA and vRNA levels to a significant extent in infected cells, a phenomenon that was not observed during transfection. NS1-AAA, in contrast, greatly enhanced the cRNA and vRNA levels and only marginally elevated the mRNA levels, thereby reconfirming that higher cytoplasmic abundance of NS1 boosts viral genome/antigenome replication during the course of infection (Figure 7B).

**Figure 7:**
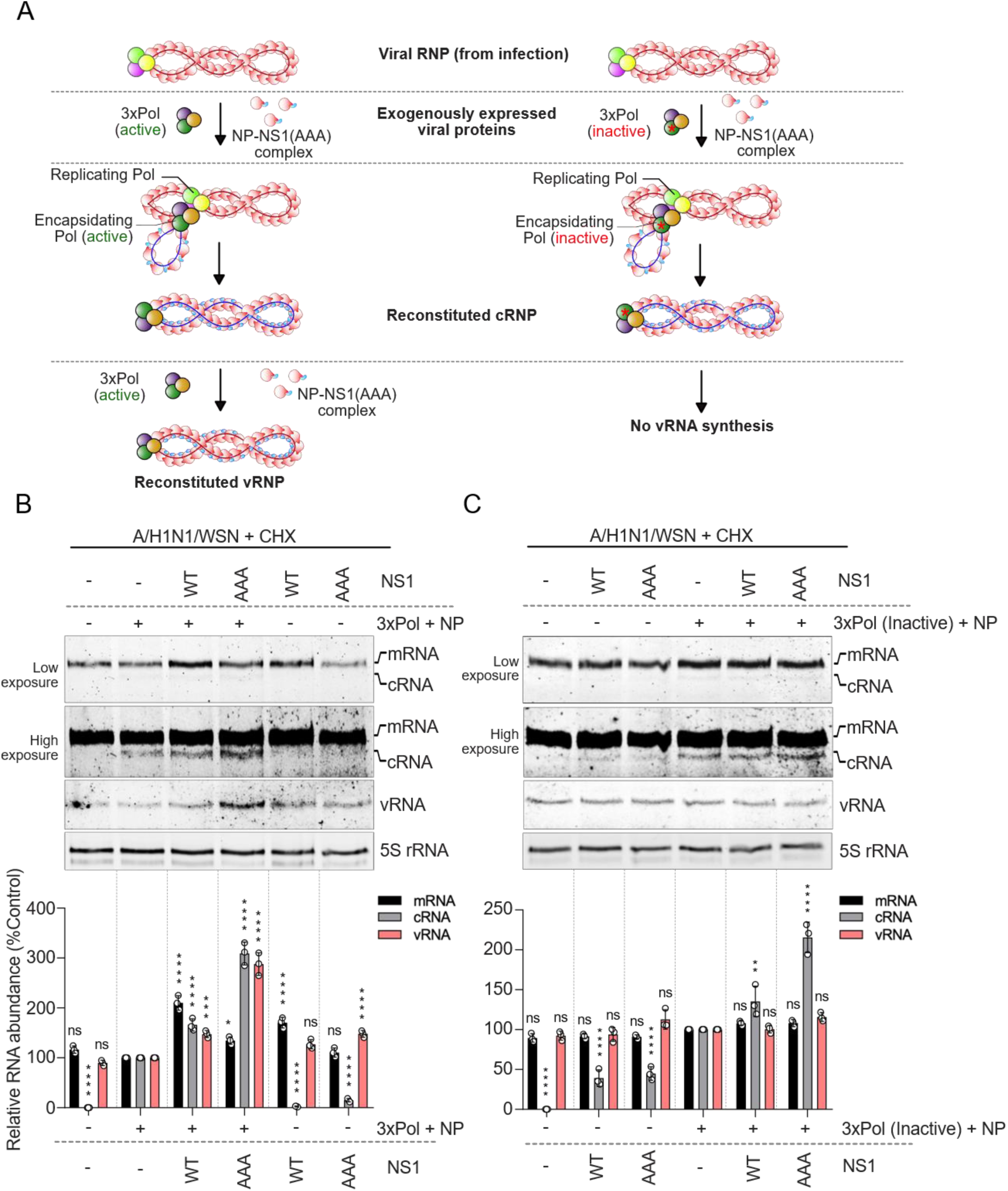
NS1 promotes v/c-RNA synthesis during influenza virus infection. (A) Diagram illustrating the experimental design for the investigation of the effect of cytoplasmic NS1 (NS1-AAA) upon viral RNA synthesis and cRNA stabilization during infection (B-C). Cells co-expressing NP with WT (B) or transcriptionally inactive RdRp (harboring PB1-D445A/D446A mutation) (C) and either NS1-WT or NS1-AAA were infected with IAV/WSN/33 (MOI 5) in the presence of cycloheximide. Abundance of viral RNA was measured by primer extension, normalized to 5S rRNA, and plotted as relative percentage of the no-NS1 control. Each experiment is performed in biological triplicate. Statistical significance was analyzed by two-way ANOVA between the individual sets with P values: ns>0.05, *P<0.05, **P<0.01, ***P<0.001, and ****P<0.0001.

The weak transcription-promoting activity of NS1-AAA might be a result of increased levels of replication and, hence, increased abundance of the vRNA template. To dissect viral transcription and replication and show that cytoplasmic NS1 specifically promotes replication by supporting the RNP assembly process, we performed cRNA stabilization assay (51). Cells expressing exogenous NP and a functionally inactive RdRp were infected, followed by cycloheximide treatment. In accordance with the previous report, a functionally inactive polymerase along with NP can support RNP assembly in trans, resulting in the production of new cRNA in the form of cRNPs (51). But these newly formed cRNPs harbor the inactive RdRp, thereby not supporting the vRNA synthesis and hence blocking subsequent rounds of mRNA synthesis. As evidenced, co-expression of NS1-AAA with NP and the defective RdRp could still support elevated levels of cRNA synthesis, which suggests (i) cytoplasmic NS1 exclusively promotes replication independent of transcription and (ii) it does so by supporting NP-dependent RNP assembly process (Figure 7C).

### NS1 orchestrates assembly of phosphorylated monomeric NP into functional RNPs

Finally, to pinpoint replication-promoting effect of NS1 in the context of the influenza virus life cycle, we employed the NP-S165D mutant influenza virus, which was reported to be severely attenuated owing to defective NP oligomerization and RNP assembly (17). The NP-S165D mutant virus replicated to 2 log lower titers compared to the WT virus, as expected (Figure 8A). Subsequently, cells overexpressing the empty vector or the NS1-AAA were infected with either wild-type (WT) A/H1N1/WSN or NP-S165D mutant viruses. Notably, the NP-S165D mutant virus showed 2-4-fold higher replication efficiency in NS1-AAA overexpressing cells compared to the control cells at 12-, 24- and 36-hours post-infection (Figures 8B and S7A). Additionally, NS1-AAA overexpression resulted in a twofold increase in WT virus replication at 12 h post-infection, although at later time points this increase was absent (Figures 8C and S7B). It is evident that higher cytoplasmic abundance of the cytoplasmic NS1 can partially rescue the virus replication defect rooted in the defective NP oligomerization and RNP assembly processes.

**Figure 8:**
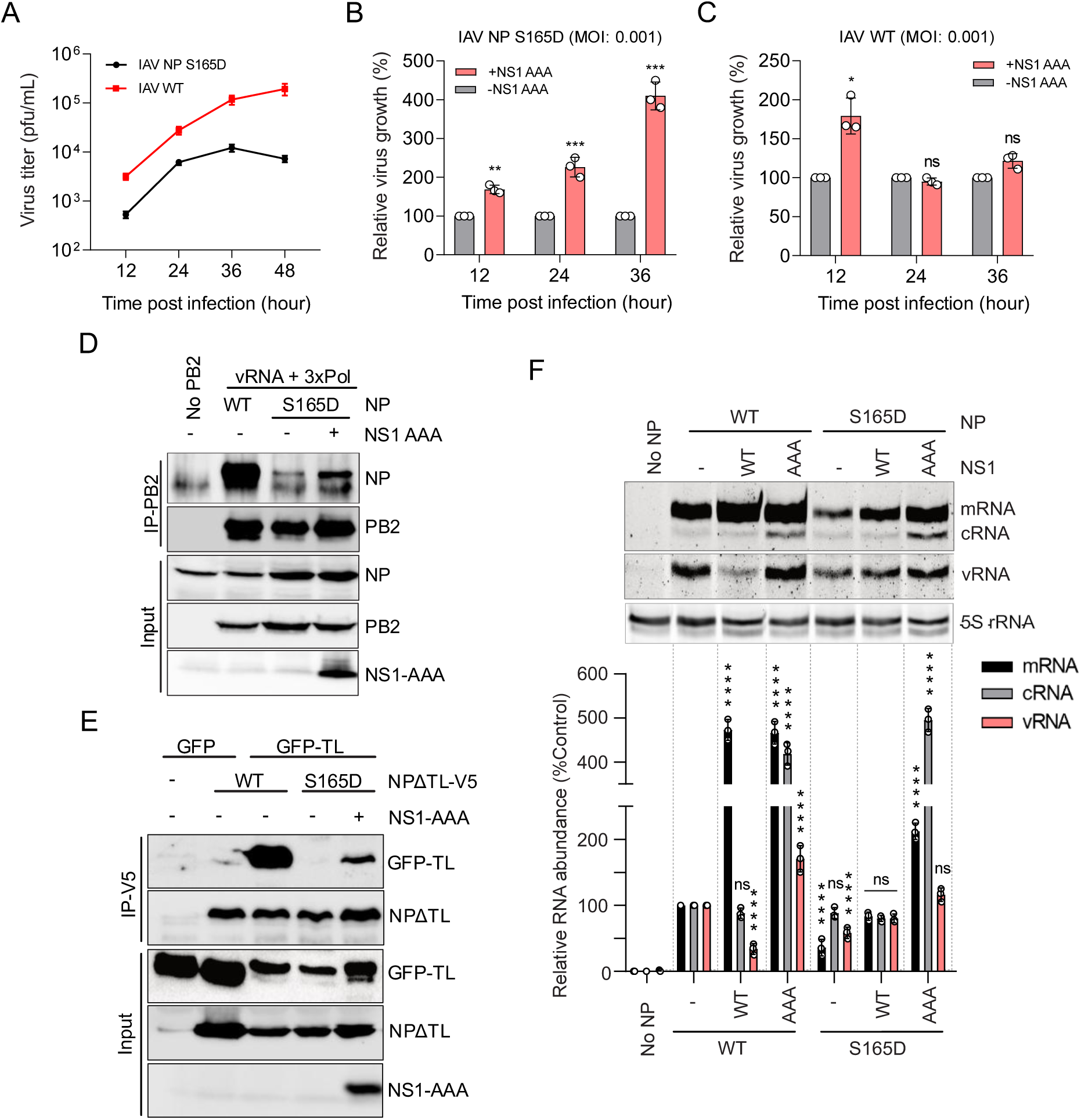
NS1 orchestrates assembly of phosphorylated monomeric NP into functional RNPs. (A) Multicycle growth of recombinant influenza A/H1N1/WSN/1933 viruses harboring either NP-WT or NP-S165D in MDCK cells at MOI 0.001. (B-C) Multicycle growth of WT (C) or NPS165D (B) virus (MOI 0.001) in control or NS1-AAA-expressing HEK293T cells. Plot representing the relative virus growth normalized to the control set (-NS1AAA) at each time point. (D) RNP reconstitution with NP WT or NP S165D in the absence or presence of NS1-AAA, followed by IP using PB2 antibody and blotting using NP antibody. (E) NPΔTL-V5 and GFP-TL were co-expressed with or without NS1-AAA and analyzed by IP using V5 antibody and blotting using GFP antibody. (F) Effects of NS1 WT or NS1-AAA on viral RNA synthesis in the context of RNPs reconstituted with NP WT or NP S165D, measured by primer extension. Each experiment is performed in biological triplicate. Statistical significance was analyzed by two-way ANOVA between the individual sets with P values: ns>0.05, *P<0.05,* *P<0.01, ***P<0.001, and ****P<0.0001.

To delve into the mechanism by which NS1-AAA may facilitate NP S165D virus replication, viral RNPs were reconstituted in cells by expressing the vRNA template, RdRp subunits (PB2-HA, PB1 and PA), and the WT or mutant NP-S165D. Large extent of WT NP was co-purified with RdRp, indicating efficient RNP formation, but the NP S165D mutant failed to do so owing to its defective oligomerization. Remarkably, in the context of NS1-AAA co-expression, NP S165D mutant supported RNP assembly to a significant extent (although lower than WT NP), thus confirming that higher cytoplasmic abundance of NS1 can drive the phospho-NP monomers (in this case, NP S165D) towards RNP assembly (Figure 8D). Notably, NS1-AAA can scaffold the tail loop groove interaction between the GFP-TL and the NPΔTL, which harbors the S165D mutation within its groove (Figure 8E & S7C). This observation provides the molecular basis of how NS1 can facilitate the oligomerization and RNP assembly of otherwise non-oligomerizing phospho-NP monomers.

Next, we investigated if the RNPs assembled with the oligomerization defective NP-S165D, aided by NS1-AAA, are functionally active in supporting viral RNA synthesis. Viral RNPs were reconstituted either with WT or mutant NP S165D in the absence or presence of WT NS1 or NS1-AAA. As expected, NP S165D, owing to its defective oligomerization, showed significant attenuation in viral RNA synthesis compared to the WT NP. Interestingly, co-expression of NS1-AAA greatly rescued the RNA synthesis defect associated with the NP S165D mutation, boosting mRNA, cRNA and vRNA levels to a large extent (Figure 8F). WT NS1, in contrast, significantly elevated mRNA synthesis and marginally increased vRNA synthesis for the S165D mutant RNPs. These data together established that cytoplasmic NS1 specifically scaffolds the monomeric NP molecules, harboring phosphorylation at their homotypic interface, to recruit them on the nascent RNP complexes, thereby facilitating RNP assembly and viral genome replication steps during the influenza virus life cycle.

## Discussion

Influenza virus RNPs are complex macromolecular machineries, central to virus replication. RNP assembly requires extensive homotypic interaction between NP molecules and their interaction with genomic or antigenomic RNAs, RdRp subunits and a number of host factors. All these interactions are required to be tightly regulated and spatiotemporally orchestrated to ensure proper and timely assembly of RNPs and hence optimum fitness of the virus. Here for the first time, we characterized NS1 as a critical regulator of RNP assembly, which initially stabilizes the newly translated and phosphorylated monomeric NP molecules within the cytoplasm and subsequently scaffolds their assembly into nascent RNPs inside the nucleus (Figure 9).

**Figure 9:**
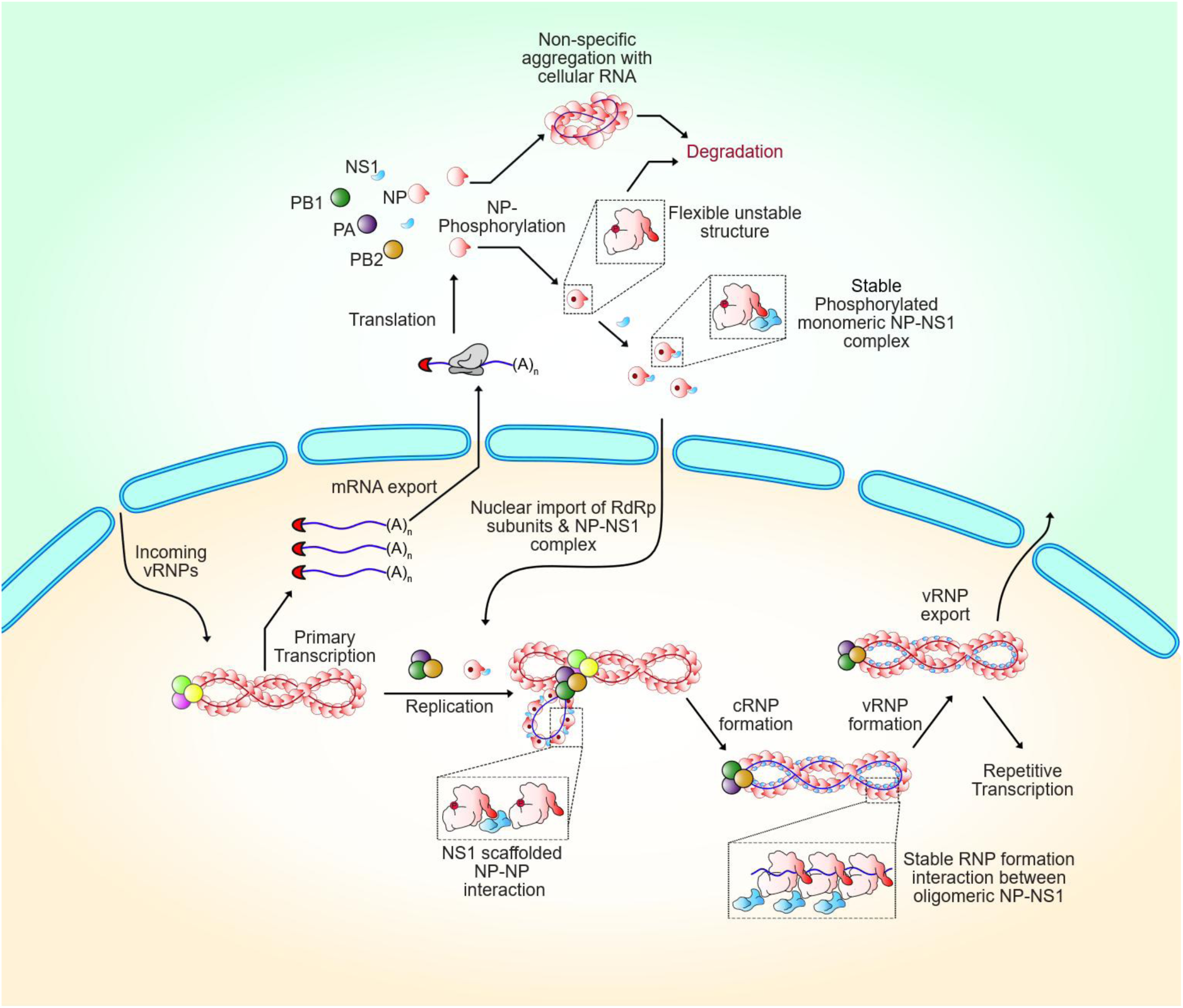
Model for NS1-mediated NP-specific chaperon activity driving NP oligomerization, RNP assembly and viral replication. Incoming RNPs execute primary transcription followed by translation of viral mRNAs, leading to the synthesis of early viral proteins, NP, PB1, PB2, PA and NS1. Newly synthesized NP monomers either can self-aggregate or get phosphorylated and kept in the monomeric form. NS1 interacts with and stabilizes the monomeric phosphorylated NPs in the cytoplasm. Once the NP-NS1 complex is translocated to the nucleus, it facilitates NP oligomerization and functional RNP assembly. During RNP assembly the NP molecules get dephosphorylated, which further stabilizes the RNP structure and increases its efficiency of RNA synthesis. Mature RNPs have NS1 still associated with them, which is facilitated through oligomeric NP-NS1 interaction.

This study presents extensive characterization of the NP-NS1 interaction in molecular detail. We showed that NS1 participates in bimodal interactions with monomeric and oligomeric NP by targeting distinct interaction interfaces. The NP-NS1 interaction is primarily governed by the NS1 RBD. In monomeric NP, the 27 amino acid long tail loop domain constitutes the major NS1-binding interface. Prior structural studies suggest that the tail loop represents a highly flexible extension in the NP structure (19, 46, 47). We utilised MD simulation to show that NS1 can specifically interact with and stabilise this flexible tail loop, thereby reducing the overall structural dynamicity and imparting stability to the monomeric NP structure. This observation is validated through cycloheximide chase experiment where co-expression of a cytoplasmic variant of NS1, NS1-AAA, significantly enhanced the in-cell stability of monomeric NP, NP S165D. Notably, WT NS1, when overexpressed in cells, majorly localize in the nucleus. In contrast, it shuttles between the nucleus and cytoplasm during early and late phases of infection, respectively. The use of NS1-AAA ensures a high abundance of NS1 in the cytoplasm, where they can capture and stabilise the NP monomers immediately after translation.

The “tail loop–groove” interaction is the major driver of NP oligomerization and its assembly into functional RNP complexes (19). Importantly, NS1’s interaction with this tail loop domain doesn’t hinder NP oligomerization, rather, it can stabilise the tail loop into an extended conformation that is suitable for interaction with the binding groove of a neighbouring protomer. In fact, NS1 can bridge the “tail loop-groove” interaction by acting as a scaffold protein. This is suggestive of the “NS1-supported NP oligomerization model,” where NS1 can bind the tail loop and stabilise the phospho-NP monomers in the cytoplasm where the concentration of NP remains low. But, within the nucleus, when the local concentration of NP rises (possibly by binding to the RNA strand), the NS1 can be displaced from the tail loop by the neighbouring NP protomers, enabling efficient NP homo-oligomer formation. Concurrently, NS1 may switch its binding preference from monomeric to oligomeric NP, promoting the formation of higher-order NP(RNA)-NS1 complexes. Interestingly, NS1 can facilitate the tail loop-groove interaction even in the presence of the phosphomimetic NP-S165D mutation within the groove that makes it sterically inaccessible for the tail loop. This data further supports the scaffolding activity of NS1, which promotes homotypic interaction between the oligomerization-defective phospho-NP molecules. This scaffolding activity of NS1 extends just beyond promoting NP oligomerization to support functional RNP assembly, enhance viral RNA synthesis, and rescue the replication defect of the NP S165D mutant virus.

Several host proteins have been previously reported to act as NP-specific chaperones (12, 13, 15, 52). RAF-2p48/BAT1/UAP56 interacts with RNA-free oligomeric NP by targeting its RNA-binding residues R174 and R175 (12, 53). UAP56, TatSF1 and Prp18 promote NP-RNA complex formation and stimulate viral RNA synthesis (12, 13, 52). Recently, ANP32A has been reported to act as a NP-specific chaperone that interacts with RNA free monomeric NP molecules to recruit them to the nascent RNA, facilitating concomitant assembly of viral RNPs during v/c-RNA synthesis (54). Notably, both ANP32 and UAP56 bind to the similar set of residues within the RNA-binding interface on NP (53, 54). These host factors predominantly localize inside the nucleus and are involved in chromatin remodeling, transcription, post-transcriptional modification, and other RNA metabolism steps. Possibly that is the reason behind influenza viruses exploiting these host factors to act as NP-specific chaperones and regulate the RNP assembly process inside the nucleus.

Non-segmented RNA viruses encode a phosphoprotein P, which acts as a Nucleocapsid protein (N)-specific chaperone thereby controlling its non-specific aggregation and RNA binding (29, 55–57). The influenza virus, instead, adopts alternative strategies; previous studies, including ours, have shown that NP phosphorylation at the homotypic interface maintains an RNA free pool of monomeric NP (16, 17). The present study elucidates an additional tier of regulation in the assembly of these phosphorylated monomeric NP molecules into functional RNP complexes. We demonstrate that the monomeric NP structure is inherently unstable and prone to degradation in the absence of binding partners. Additionally, NP monomers might be more susceptible to ubiquitination by host E3 ligases, including TRIM family proteins, leading to proteasomal degradation (58–60). NS1 imparts structural compactness by binding to these phosphorylated monomeric NPs thereby enhancing their in-cell stability. Eventually, these NP-NS1 complexes are trafficked into the nucleus and recruited on the nascent RNA chain concomitant to its synthesis. Here, NS1 acts as a scaffold to facilitate the assembly of the phospho-NP monomers into the pre-mature RNPs which eventually oligomerize and form stable RNP complexes. NS1 may coordinate with ANP32, UAP56 and other host chaperones and synergistically regulate NP oligomerization, RNA binding and ultimately the assembly of functional RNP complexes. It would also be interesting to investigate how NP-NS1 interaction can influence NP’s interaction with host kinases that are reported to associate with RNPs and the export machinery that is instrumental for nuclear-cytoplasmic transport of the newly formed RNPs (18, 61, 62).

Together, our work elucidates a completely novel role of NS1 proteins in regulating the assembly of influenza virus replication machinery through characterization of the NP-NS1 interaction in molecular detail. This protein-protein interaction interface may serve as a novel target for the development of next-generation antivirals against influenza viruses.

## Materials and methods

### Cells, viruses, and antibodies

Human embryonic kidney (HEK) 293T cells and Madin Darby canine kidney (MDCK) were maintained in Dulbecco’s modified Eagle’s medium (DMEM, Gibco) supplemented with 10% heat-inactivated fetal bovine serum (FBS, Gibco), 100 µg/mL penicillin-streptomycin (Gibco), and 2 mM L-glutamine (Gibco). Human lung epithelial cells (A549) were maintained in Dulbecco’s modified Eagle’s medium: Nutrient Mixture F-12 (DMEM/F-12, Gibco) supplemented with 10% heat-inactivated fetal bovine serum (FBS, Gibco), 100 µg/mL penicillin-streptomycin, 1X MEM Non-Essential Amino Acids (NEAA, Gibco), 1 mM sodium pyruvate, and 2 mM l-glutamine. All cells were maintained at 37°C with 5% CO_2._ Influenza A virus strain, A/H1N1/Wilson-Smith/1933 (H1N1/WSN/1933), encoding PB2 protein fused to a C-terminal Flag tag, was used for infection studies (63).

Antibodies: anti-HA (C29F4, Cell Signaling Technology, CST), anti-V5 (D3H8Q, CST), anti-NP (H16-L10-4R5, BEI Resources), anti-FLAG M2 (F1804-1MG, Sigma), anti-influenza A virus RNP (NR-3133, BEI Resources), IAV NS1 antibody (PA5-32243, Invitrogen), anti-GFP (BB-AB006, Bio-Bharati), anti-GAPDH (BB-AB0060, Bio-Bharati), anti-H3 (BB-AB0055, Bio-Bharati), rabbit anti-mouse IgG (A9044, Sigma), and goat anti-rabbit IgG (12-348, Sigma) antibodies were used as per manufacturer protocol.

### Plasmids

The plasmids pCDNA3-PB2-3XFLAG, pCDNA3-PA, and pCDNA3-PB1, along with pCDNA6.2-NP-V5, pEGFP-NP, and pCDNA3-NS1-3XFLAG, were utilized for the expression of influenza A/WSN/33 (H1N1) viral proteins. The plasmid pET28a-NΔ7NP was employed for the bacterial expression of wild-type A/WSN/1933 (H1N1) nucleoprotein (NP) carrying a seven-amino acid deletion at the N-terminus and a C-terminal His tag, as previously described (17). The constructs pET28a-NS1 RBD and pET28a-NS1 FL were designed to express the RNA-binding domain (RBD) and full-length (FL) NS1 proteins of A/WSN/1933 (H1N1), respectively. The reporter plasmids pHH21-vNA-luc and pHH21-vHA-luc encode firefly luciferase in a negative-sense orientation, flanked by the untranslated regions (UTRs) of the NA and HA gene segments of A/WSN/1933 (H1N1) respectively, were generated based upon the protocol described previously (64). Truncated NS1 constructs (RBD and ΔRBD) were cloned into the pCDNA3-3XFLAG vector for mammalian expression. The pEGFP-TL construct was created as previously discussed (17). Recombinant viruses were produced utilizing a bidirectional reverse genetics plasmid approach, as previously described (65–67). All constructs were verified by Sanger sequencing.

### Site-directed mutagenesis

Site-directed mutagenesis was conducted on the NP bacterial expression construct to produce pET28a-NΔ7NP E339A, S486A, S165D, and ΔTL variants and on the NP mammalian expression construct to generate pCDNA6.2-NP E339A-V5, S486A-V5, S165D-V5, and ΔTL-V5 mutants. NS1 AAA (R148A/E152A/E153A) was generated similarly. Bacterial NS1 RBD and full-length NS1 constructs carrying R38A/K41A substitutions were generated similarly. Mutagenesis was performed through PCR amplification of the parental plasmids using PfuTurbo DNA polymerase (Agilent Technologies), followed by DpnI (NEB) digestion and transformation into competent *E. coli* DH5α cells. All substitutions were confirmed using Sanger sequencing.

### Recombinant protein expression and purification

Wild type & mutants of NP and full-length NS1 constructed in the pET28a vector were expressed in *E. coli* Rosetta DE3 in LB medium containing 50 µg/ml Kanamycin. NS1 RBD constructed in pRSF vector was expressed in *E. coli* strain BL21 DE3 in LB medium containing 50 µg/ml Kanamycin. Protein expression was induced by 0.5 mM IPTG for 18-20 hr. at 16 °C. Cells were harvested and resuspended in lysis buffer consisting of 50 mM Tris-HCl, pH 7.5, 300 mM NaCl, and 1 mM DTT, followed by sonication on ice. The clarified lysates were loaded onto Ni-NTA agarose (Bio-Rad Profinity). To remove RNA associated with NP and NS1, the resin was washed with RNase A and then with high-salt buffer prepared in lysis buffer supplemented with 1.5 M NaCl. Bound proteins were eluted with imidazole-containing buffer and further purified by size-exclusion chromatography on ENrich SEC650 column (Bio-Rad) or Superdex 200 Increase 10/300 GL column (GE Healthcare) equilibrated in 50 mM Tris-HCl, pH 7.5, 150 mM NaCl, and 1 mM DTT.

### His-tag Co-elution assay

To assess NP–NS1 interactions in vitro, purified untagged wild-type or mutant NPs were incubated with 6×His-tagged NS1 RBD in binding buffer containing 50 mM Tris-HCl, pH 7.5, 150 mM NaCl, 1 mM DTT, and 5 mM imidazole at 4°C overnight. The mixture was then incubated with Ni-NTA agarose for 1 h at 4°C. The flow-through was collected, and the resin was washed sequentially with buffer containing 10 mM and 50 mM imidazole. Bound proteins were eluted with 250 mM imidazole, resolved by 15% SDS-PAGE, and visualized by Coomassie Brilliant Blue staining.

### Surface plasmon resonance

SPR analysis was performed on a Biacore T200 at 25°C. NP WT or mutant proteins were immobilized on a Series S CM5 sensor chip by amine coupling using HBS-EP buffer (10 mM HEPES, pH 7.4, 150 mM NaCl, 3 mM EDTA, 0.005% P20 (v/v)). NS1 RBD or full-length NS1 was injected at a flow rate of 30 µL/min for 60 or 90 s, respectively, using a running buffer containing 50 mM Tris-HCl, pH 7.5, 200 mM NaCl, and 0.5 mM DTT. For reciprocal binding assays, NS1 was immobilized on the CM5 chip and NP mutants (NP E339A, NP S486A and NP delTL) were injected at increasing concentrations for 150 s. The chip surface was regenerated with 10 mM glycine, pH 2.5, after each cycle. For each experimental set, double referencing was performed using blank immobilized surfaces and buffer injections to eliminate nonspecific and bulk contributions.

### Analytical size exclusion chromatography

For complex formation analysis, purified NP proteins (NP E339A or NP S486A) and NS1 RBD were mixed in a 1:1 molar ratio and incubated overnight at 4°C. The complex was resolved on Superdex® 200 Increase 10/300 GL column equilibrated in 50 mM Tris-HCl pH 7.5; 150 mM NaCl and 1 mM DTT.

### Transfection, co-immunoprecipitation study & western blotting

HEK 293T cells were transfected with the indicated plasmids using lipofectamine 3000 (Invitrogen) into 1.5X10^6^ cells. The transfected cells were lysed in radio-immunoprecipitation assay (RIPA) buffer (50 mM Tris-HCl pH 7.5, 150 mM NaCl, 2 mM EDTA, 1% NP-40, 0.5% deoxycholate, 0.1% SDS), and crude protein was collected after centrifugation. RNP reconstitution was performed by transfecting 3X polymerase (PB2, PB1 and PA) and NP with or without NS1 AAA. The transfected cells were lysed in radio-immunoprecipitation assay (RIPA-DOC) buffer (50 mM Tris-HCl pH 7.5, 150 mM NaCl, 2 mM EDTA, 1% NP-40, 0.1% SDS), and crude protein was collected after centrifugation as described (17). Lysates were pre-cleared using protein-A agarose beads (BioBharti) in 5 mg/ml BSA containing RIPA buffer to reduce nonspecific binding. The samples were then incubated with appropriate antibodies and captured on Protein A Dynabeads (Invitrogen) or anti-FLAG beads (Sigma). Beads were subsequently washed with RIPA buffer containing 500 mM NaCl once and with regular RIPA buffer twice before elution and western blot analysis.

For the NP oligomerization assay, NP-V5 and NP-GFP were co-expressed in the presence of increasing concentrations of NS1AAA in HEK 293T cells. The transfected cells were lysed in co-immunoprecipitation (Co-IP) buffer (50 mM Tris-HCl pH 7.5, 150 mM NaCl, and 1% NP-40). Lysates were pre-cleared as discussed above. The samples were then incubated with appropriate antibodies in the presence of RNaseA (Thermo Scientific) at 4⁰C overnight. Finally, the immunocomplexes were captured on Protein A Dynabeads. The washing step was performed as per the protocol mentioned above.

### Polymerase activity assay

HEK293T cells were transfected using lipofectamine 3000 (Invitrogen) in triplicate with plasmid pCDNA3-PA, pCDNA3-PB1, pCDNA3-PB2, pCDNA6.2-NP-V5, pHH21-vNA-LUC or pHH21-vHA-LUC viral reporter constructs (bearing 5’ and 3’-UTRs corresponding to the NA and HA segments respectively), PCNDA3-NS1-3xFLAG (WT or mutants), and pRL-SV40 reporter construct as transfection control. Cells were lysed at 24 hr. post-transfection, and luciferase activity was measured by using the Dual-Luciferase® Reporter Assay System (Promega) in the GloMax 20/20 instrument (Promega).

### Primer extension analysis

HEK293T cells were transfected using lipofectamine 3000 (Invitrogen) with plasmids pCDNA3-PA, pCDNA3-PB1, pCDNA3-PB2, pCDNA6.2-NP-V5 and pBD-NP (harboring a premature stop codon in the NP ORF) to reconstitute full-length RNP or with pCDNA3-PA, pCDNA3-PB1, pCDNA3-PB2 and NP77 (49) (harboring NP segment specific-5’ and 3’ UTRs only) to reconstitute NP-free mini RNPs, in the absence or presence of PCNDA3-NS1-3xFLAG (WT and mutants). Cells were lysed at 24 hr. post-transfection, and total cellular RNA was extracted using Trizol reagent (Invitrogen). Primer extension was conducted using fluorescently labeled NP segment-specific primers for vRNA, mRNA, and 5S rRNA as described earlier (48, 49, 62). The levels of different RNAs were determined by running urea-PAGE and visualized through Chemidoc MP (Bio-Rad).

### cRNA stabilization assay

The cRNA stabilization assay was performed as originally described by Vreedy et al. (51, 68). HEK293T cells were transfected with plasmids encoding PA, PB1-D445A/D446A, PB2-FLAG, NP-V5 in the absence or presence of plasmid encoding NS1 or NS1AAA using lipofectamine 3000 (Invitrogen). After 24 h.p.t., cells were infected with the IAV/WSN/1933 virus at MOI 5 and subsequently treated with 1 mM Cycloheximide (Sigma). Infected cells were harvested after 6 h.p.t., and total cellular RNA was extracted using Trizol reagent (Invitrogen). The primer extension analysis was conducted as per the protocol mentioned earlier.

### Infection

A549 cells were infected with the IAV/WSN/1933 virus at MOI 0.1 and harvested at 24 hours post-infection. Viral NP proteins were immunoprecipitated using anti-NP antibodies, and co-immunoprecipitated NS1 was detected by Western blotting using anti-NS1 antibodies. For immunofluorescence analysis, A549 cells were infected with IAV/WSN/33 virus at MOI 5. at different times (2, 4 and 6 hours) post infection, cells were fixed, blocked, and stained for appropriate antibodies. For fractionation study, A549 cells were infected with IAV/WSN/33 virus at MOI 2, and infected cells were harvested 6 hours post infection.

### Immunofluorescence

A549 cells were transfected with NS1 or NS1-AAA individually for 24 hours. Following transfection and infection, cells were fixed with 3% formaldehyde, quenched with glycine, permeabilized with 0.1% Triton X-100 in PBS, and blocked with 3% BSA. Cells were then incubated with the appropriate primary and fluorescent secondary antibodies, counterstained with DAPI, and imaged by fluorescence microscopy (Leica). Nuclear-to-total fluorescence intensity ratios were quantified in ImageJ from at least 50 randomly selected cells per condition across three independent biological replicates.

### Sub-cellular Fractionation

Infected or transfected cells were washed with cold 1X PBS once and lysed with cytoplasmic lysis buffer (10vmM Tris pH 7.5, 10vmM KCl, 1.5vmM MgCl_2,_ 0.2vmM EDTA, 0.05% Triton X-100, 1 mM DTT and 1X protease and phosphatase inhibitor) for 10 mins on ice. Then cells were centrifuged at 200xg for 3 mins. The supernatant was collected as cytoplasmic fraction. The pellet was washed and lysed in RIPA buffer for 30 mins on ice. Both fractions were centrifuged at 21, 000Xg for 10 mins to remove debris and subjected to SDS-PAGE and Western blotting. GAPDH and Histone H3 were used to track the cytoplasmic and nuclear fractions respectively.

### Cycloheximide chase assay

HEK 293T cells were transfected with NP S165D alone or together with NS1 AAA and treated with 1 mM cycloheximide at 12 h post-transfection (Sigma). Harvested after 3 or 6 h, followed by western blot analysis.

### Generation of recombinant viruses and multicycle replication assay

Recombinant WT and NP mutant viruses were rescued using the influenza bidirectional reverse genetics system by co-transfecting HEK 293T/MDCK co-cultures with virus rescue plasmids, pBD PB2, pBD* PB1, pBD PA, pBD NP or pBD NP S165D, and pTMΔRNP using Lipofectamine 3000 reagent (Invitrogen) as described previously (48, 62). The media was replaced with virus growth media (VGM: 1× DMEM, 1× PenStrep, 0.2% bovine serum albumin [BSA], 25 mM HEPES buffer, 4 mM GlutaMax and 0.5 μg/mL L-1-tosylamido-2-phenylethyl chloromethyl ketone [TPCK]-treated trypsin) after 24 hours post transfection. Supernatants were collected after 36–48 h, amplified in MDCK cells for two passages, and titrated by plaque assay.

HEK293T cells were transfected with plasmid encoding NS1AAA. 24 hours post-infection, cells were infected with wild-type A/WSN/1933 (H1N1) virus or viruses harbouring the NP S165D mutation at MOI of 0.001 in three biological replicates. Supernatants were collected at 12-, 24- and 36-hour post-infection and subsequently titrated by plaque assay.

### Molecular docking

Protein–protein docking was performed with the NP monomer extracted from the NP trimeric assembly (PDB ID: 2IQH) and the NS1 monomer derived from the dimeric protein (PDB ID: 4OPH) (19, 69). Reverse mutations were introduced in silico at NS1 positions 38 and 41 to reconstruct the wild-type sequence. Rigid-body protein–protein docking was then carried out using HADDOCK2.4 (70). The NP tail-loop and NS1 residues 38–42 was defined as the interaction interface. The top-ranked docked complex was selected for subsequent computational analyses.

### Molecular Dynamics Simulation

All-atom molecular dynamics (MD) simulations were conducted using GROMACS 2023. 3 in conjunction with the CHARMM27 force field (71). To refine the relative structural orientation of the NP–NS1 complex (obtained through molecular docking) and obtain a more biologically relevant interfacial conformation, an initial short MD simulation of 50 ns was first carried out for the docked NP–NS1 complex. A representative structure selected from the dominant cluster obtained through cluster analysis of this trajectory was subsequently used as the starting configuration for a longer production simulation for the NP-NS1 complex. 200 ns production simulations were performed for the NP-monomer, the NP-NS1 heterodimer, and the NP-NP homodimer, with the latter serving as a comparative reference for interchain binding affinities. All systems were solvated using the TIP3P water model and neutralized with Na^+^ and Cl^-^ ions. Following energy minimization via the steepest descent algorithm, each system was equilibrated for 100 ps under the NVT ensemble and 500 ps under the NPT ensemble. Long-range electrostatic interactions were treated using the Particle Mesh Ewald (PME) method (72). Production MD was executed for 200 ns with a 2-fs time step, maintaining a constant temperature of 300 K via the V-rescale thermostat and a pressure of 1 atm using the Parrinello-Rahman barostat (73, 74).

### MM-PBSA Calculations

Binding free energies for the NP–NS1 heterodimer and NP–NP homodimer were estimated using the gmx_MMPBSA (v1.6.4) package (75, 76). A total of 50 frames, extracted from the last 50 ns of the equilibrated MD trajectories, were employed for the analysis. The binding free energy (ΔG_bind)_ of the protein complexes was computed according to: ΔG_bind =_ G_peptide-complex –_ (G_peptide-1 +_ G_peptide-2)_

Where G_peptide-complex r_epresents the free energy of the bound protein dimer (NP-NS1 or NP-NP), and G_peptide-1 a_nd G_peptide-2 c_orrespond to the free energies of the constituent protein chains in their unbound states. To account for the entropic contribution, the Interaction Entropy (IE) approximation was applied to arrive at the final ΔG_bind._

### Statistical analysis

All experiments were executed with at least three independent biological replicates, each carried out in triplicate (n = 3). Data are expressed as mean ± standard deviation (SD). Graphs were generated utilizing GraphPad Prism (version 10.1.2). Comparisons between the two groups were performed using a two-tailed unpaired Student’s t-test. For comparisons across multiple groups, one-way or two-way analysis of variance (ANOVA) was applied as suitable to the experimental design. Densitometric analysis was conducted with Bio-Rad Image Lab software.

Statistical significance was defined at P < 0.05. Significance levels are denoted as follows: ns (not significant, P > 0.05), * (P ≤ 0.05), ** (P ≤ 0.01), *** (P ≤ 0.001), and **** (P ≤ 0.0001).

## Acknowledgments

We sincerely acknowledge Andrew Mehle (University of Wisconsin Madison) for providing the plasmid pEGFP-TL. This work was primarily supported by the Core Research Grant (CRG) from the Anusandhan National Research Foundation (ANRF)/Science and Engineering Research Board, Department of Science and Technology (SERB-DST) [CRG/2022/003628]. A.M. acknowledges Indian Council of Medical Research (ICMR) Small Grant [IIRPSG-2025-01-00024] and the Department of Biotechnology (DBT), Ministry of Science & Technology, Government of India [BT/PR58658/BMS2/156/208/2025] for providing additional financial support. S.D. acknowledges the University Grants Commission (UGC), Government of India, for providing doctoral fellowship support. A.D. acknowledges the Ministry of Education (MoE), Government of India, for the Prime Minister’s Research Fellowship (PMRF). The authors also acknowledge the Central Research Facility (CRF), Indian Institute of Technology Kharagpur, for providing access to essential instrumentation and research infrastructure. The funding agencies had no role in the design of the study, data collection and interpretation, or the decision to submit the work for publication.

