## Supplementary figures for "Influenza virus Non Structural protein 1 (NS1) Chaperones Nucleoprotein (NP) Oligomerization to Coordinate Ribonucleoprotein Assembly and Genome Replication"

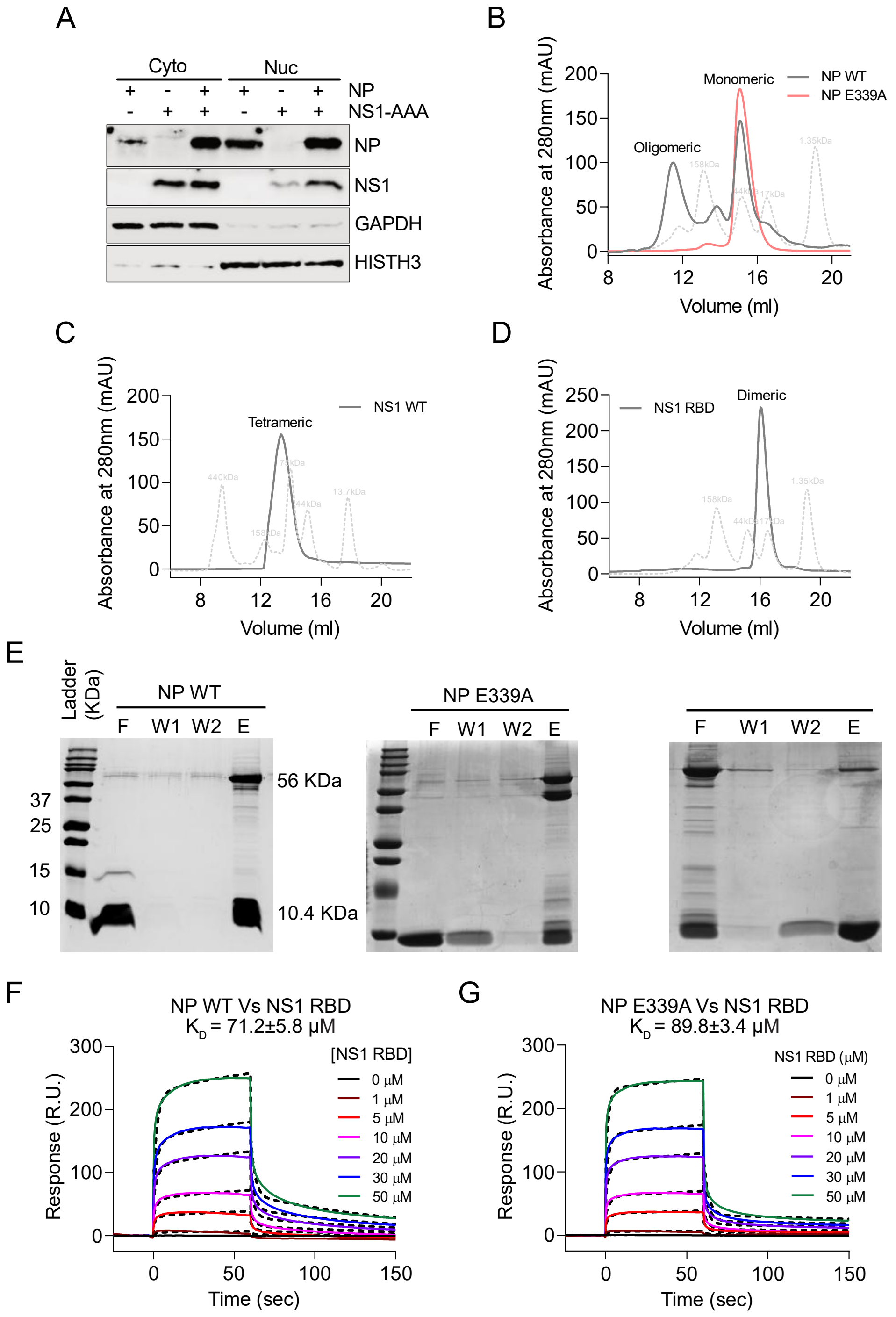


**Figure S1: NP and NS1 proteins engage in direct interaction in both nuclear and cytoplasmic compartments.** (A) V5-tagged NP was co-expressed with FLAG-tagged NS1 WT or NS1-AAA in A549 cells, fractionated, and analyzed by immunoblotting. GAPDH and Histone H3 were used to track cytoplasmic and nuclear fractions. (B-D) Bacterial-expressed recombinant NP WT (B), NP E339A (C), NS1 WT and NS1 RBD (D) proteins were analyzed by size exclusion chromatography. Oligomeric and monomeric peaks were marked. (E) His tag co-elution assay using histidine-tagged NS1-RBD and untagged NP proteins (NP WT, NP E339A and NP S486A). Eluted fractions were separated using 15% SDS-PAGE and observed using Coomassie Brilliant Blue staining. W1 = 10 mM wash fraction, W2 = 50 mM wash fraction, and E = eluted fraction. (F-G) Surface plasmon resonance study was performed by immobilizing NP WT (F) and NP E339A (G) proteins on CM5 chip, and different concentrations of NS1 RBD were run over it. Sensogram of the interaction was fitted, and the $K_D$ of the respective interaction was determined through the Biacore instrument software.

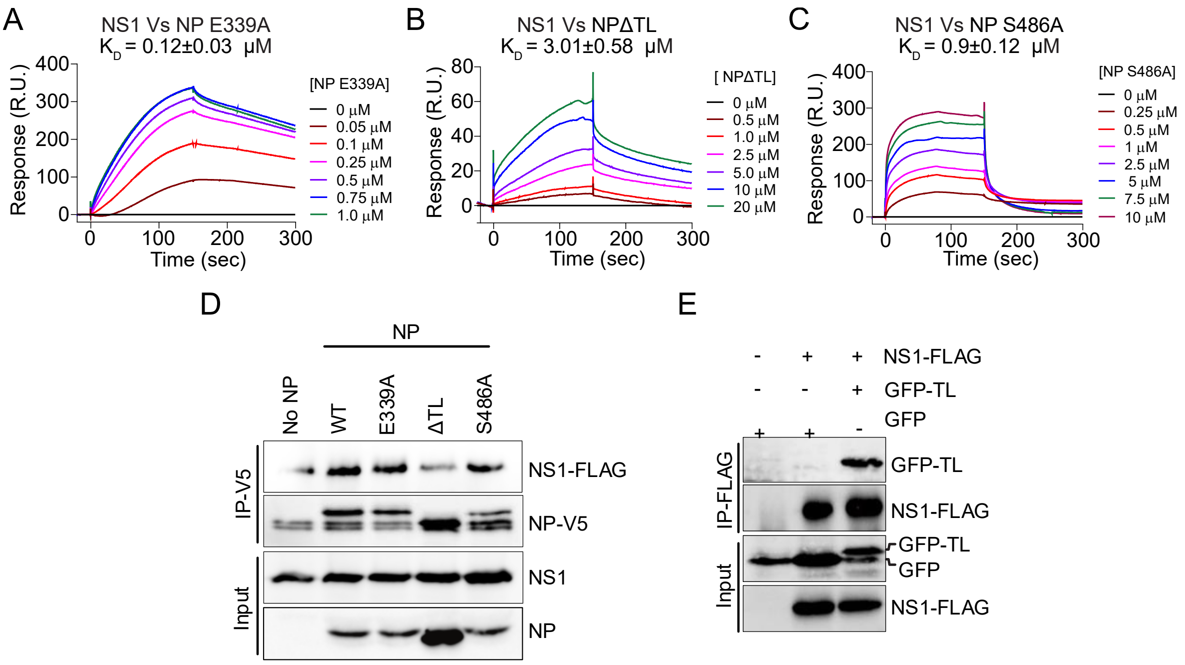


**Figure S2: Bimodal nature of NP-NS1 interaction.** Sensogram of surface plasmon resonance study between NS1 protein and NP mutants E339A (A), ΔTL (B) and S486A (C). (D) V5-tagged NPs WT or mutants (E339A, ΔTL & S486A) were co-expressed with FLAG-NS1 in HEK293T cells and analyzed by co-IP in the presence of RNaseA. (E) GFP-TL was co-expressed with NS1 WT in HEK293T cells and analyzed by co-IP using anti-FLAG beads.

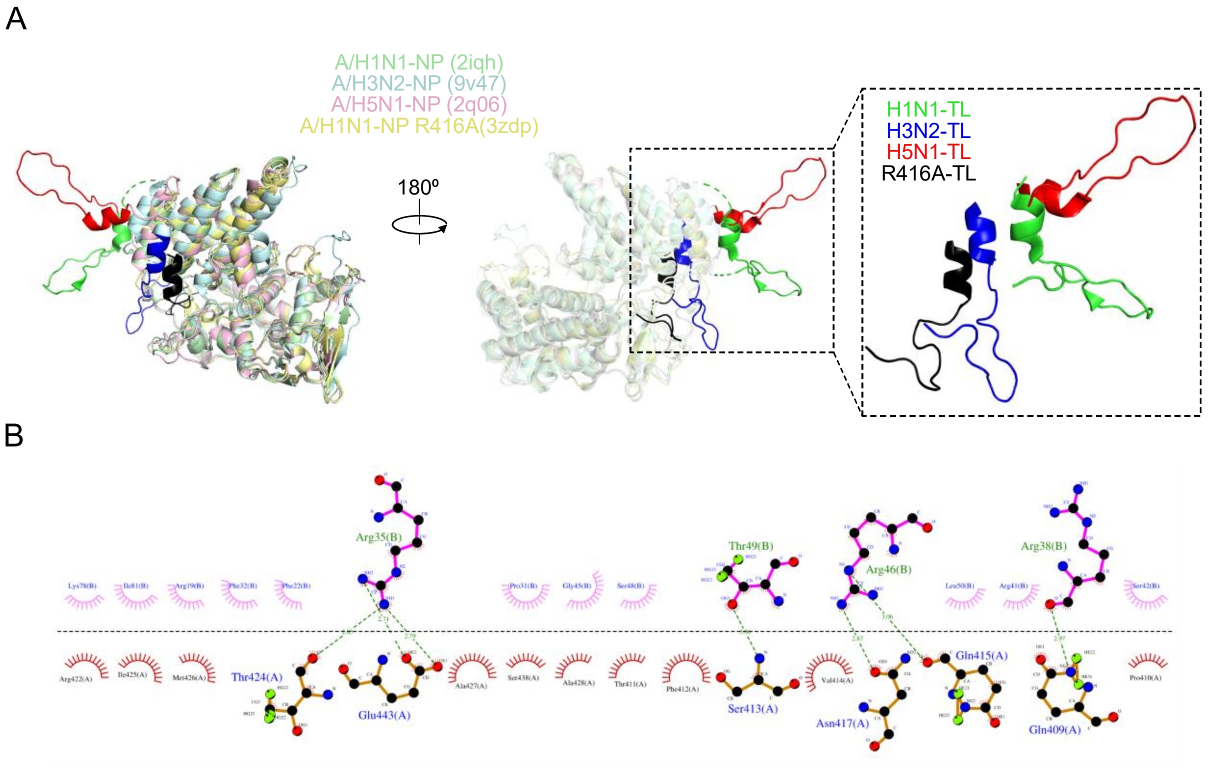


**Figure S3: NS1 stabilizes NP monomer structure by reducing flexibility of the tail loop domain.** (A) Structural superimposition of NPs from different Influenza A virus strains (H1N1, H3N2 and H5N1), highlighting the tail loop domain of each NP structure. H1N1-TL in green, H3N2-TL in blue, H5N1-TL in red and H1N1-NP R416A in black. (B) LigPlot analysis of this lowest-energy conformation of the NP-NS1 complex highlights the key interfacial residue contacts.

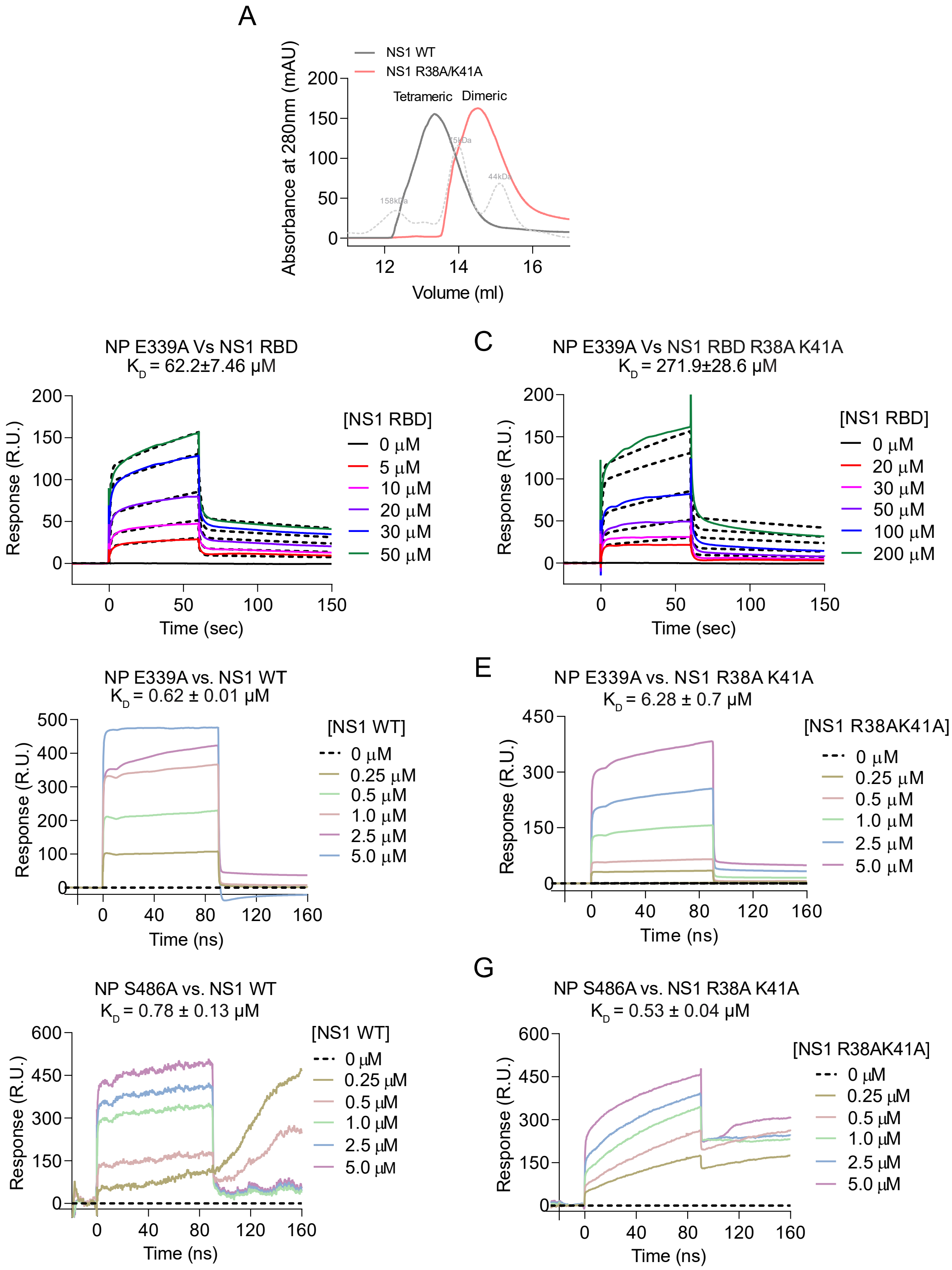


**Figure S4: NS1 acts as a cytoplasmic chaperone to stabilize monomeric NP.** (A) Size exclusion chromatography profile of recombinant purified NS1 WT and NS1-R38A/K41A resolved through Superdex® 200 Increase 10/300 GL. (B-C) Surface plasmon resonance study to measure the interaction affinity between NP E339A and NS1 RBD (B) or NS1 RBD-R38A/K41A (C). KD of the respective interaction was determined after fitting. (D-G) Sensogram of surface plasmon resonance study between NS1 WT and NS1 R38A/K41A with NP E339A or NP S486A.

A

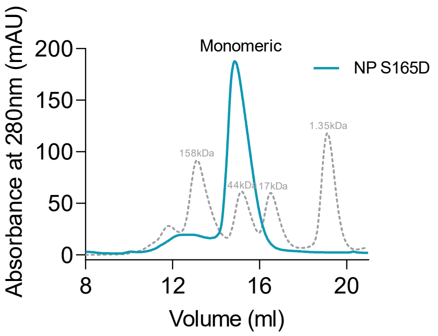


**Figure S5:** (A) Size exclusion chromatography profile of recombinant purified NP S165D resolved through ENrich™ SEC650 column.

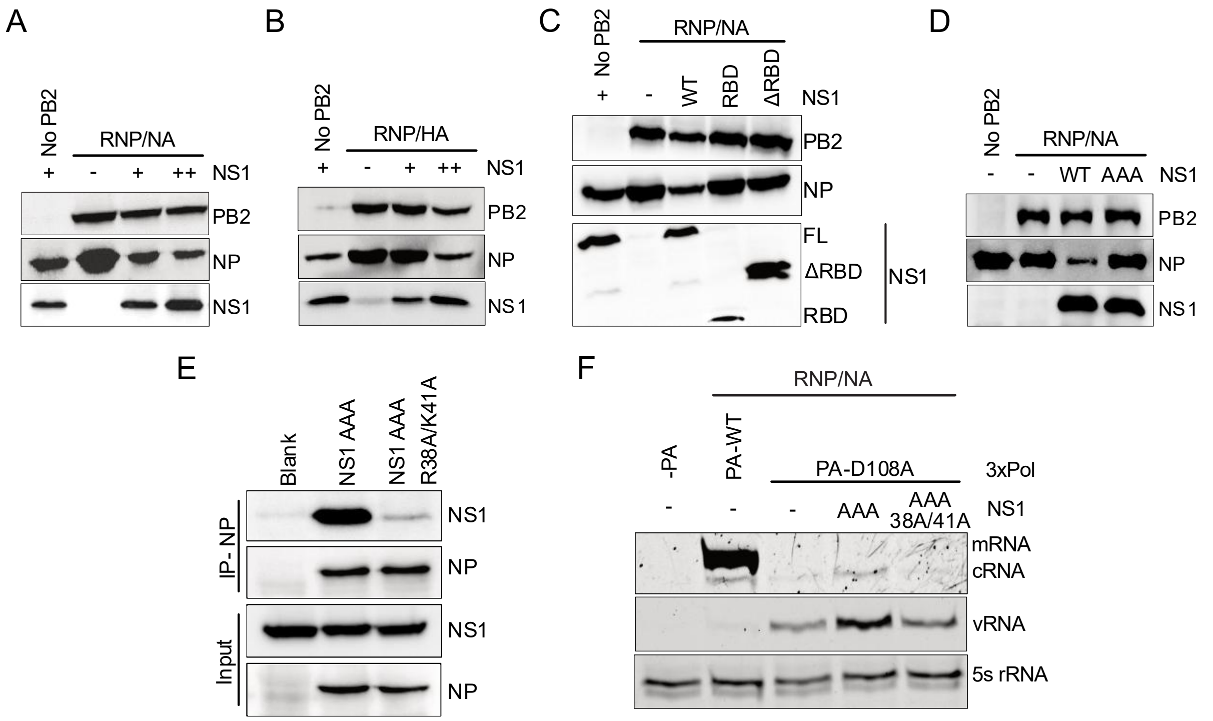


**Figure S6: Cytoplasmic NS1 promotes viral genome replication in NP dependent manner.** (A-D) Western blotting analysis of polymerase activity assay. (E) Co-immunoprecipitation of NP-V5 and NS1-AAA-R38A/K41A in HEK293T cells according to the protocol depicted in Fig. 1D. (F) Effect of NS1-AAA R38A/K41A on the viral RNA synthesis in the transcription-defective (PA-D108A) RNP/NP system examined through primer extension analysis.

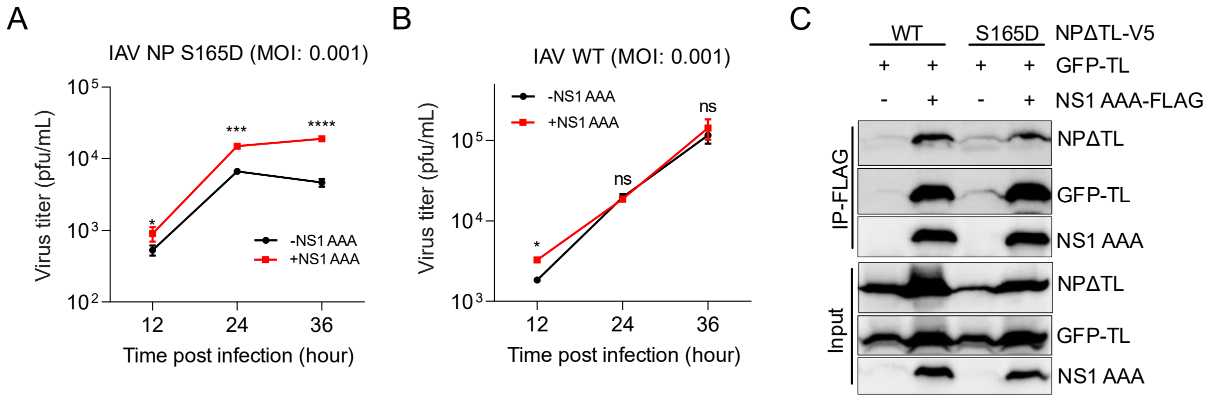


**Figure S7: NS1 orchestrates assembly of phosphorylated monomeric NP into functional RNPs.** (A & B) Multicycle replication growth curve of WT (B) and NP S165D (A) mutant virus (MOI: 0.001) in the absence or presence of NS1-AAA. Plot representing the viral titer at each time point. (C) V5-tagged NP delTL or NP delTL S165D and GFP-TL plasmids were co-transfected with NS1-FLAG, and immunoprecipitation was done using the similar procedure as mentioned in 5F.
